# The Forget-Me-Not dHCP study: 7 Tesla high resolution diffusion imaging in the unfixed post-mortem neonatal brain

**DOI:** 10.1101/2021.06.24.449666

**Authors:** Wenchuan Wu, Luke Baxter, Sebastian W Rieger, Eleri Adams, Jesper LR Andersson, Maria Cobo Andrade, Foteini Andritsou, Matteo Bastiani, Ria Evans Fry, Robert Frost, Sean Fitzgibbon, Sean Foxley, Darren Fowler, Chris Gallagher, Amy FD Howard, Joseph V Hajnal, Fiona Moultrie, Vaneesha Monk, David Andrew Porter, Daniel Papp, Anthony Price, Jerome Sallet, Michael Sanders, Dominic Wilkinson, Stephen M Smith, Rebeccah Slater, Karla L Miller

## Abstract

Diffusion MRI of the neonatal brain allows investigation of the organisational structure of maturing fibres during brain development. Post-mortem imaging has the potential to achieve high resolution by using long scan times, enabling precise assessment of small structures. The Forget-Me-Not study, part of the Developing Human Connectome Project (dHCP), aims to acquire and publicly distribute high-resolution diffusion MRI data for unfixed post-mortem neonatal brain at 7T with a custom-built head coil. This paper describes how the study addressed logistical, technical and ethical challenges relating to recruitment pipeline, care pathway, tissue preservation, scan setup and protocol optimisation. Results from the first subject recruited to the study demonstrate high-quality diffusion MRI data. Preliminary voxel-wise and tractography-based analyses are presented for the cortical plate, subplate and white matter pathways, with comparison to age-matched in vivo dHCP data. These results demonstrate that high quality post-mortem data can be acquired and provide a sensitive means to explore the developing human brain, as well as altered diffusion properties consistent with post-mortem changes, at high resolution.

## Introduction

Advances in infant magnetic resonance imaging (MRI) acquisition and analysis have improved our understanding of how the human brain develops in early life (Counsell et al., 2019). Further, such data have provided insight into how premature birth and early brain injury impact structural brain development (Dubois et al., 2014; Miller et al., 2005). Over the past decade, diffusion MRI (dMRI) in particular has arisen as a powerful method for studying the development of structural connectivity in the brain (Qiu et al., 2015). This has motivated major imaging projects such as the Developing Human Connectome Project (dHCP) (http://www.developingconnectome.org), which aims to provide comprehensive mapping of connectivity patterns in the infant brain.

Compared to in vivo image acquisition, post-mortem brain imaging allows for acquisition of higher spatial resolution data with improved tissue contrast, facilitating the link between MRI imaging and gold-standard histological findings (Roebroeck et al., 2019). Particularly relevant to the infant population, post-mortem MRI images are not contaminated with head motion and physiological signal artefacts, which are often prominent detrimental features of infant MRI data acquired in vivo (Bastiani et al., 2019) that can significantly confound biological interpretation (Yendiki et al., 2014)

Post-mortem MRI of prematurely-born infants who died shortly after birth and of fetuses following miscarriage or termination has allowed for comprehensive regional assessment of volumetric growth trajectories (Kinoshita et al., 2001), resolution of superficial and deep subplate layers (Pogledic et al., 2020), and delineation of the organisational structure of maturing connectional fibres (Huang et al., 2009; Kolasinski et al., 2012; Takahashi et al., 2012). Additionally, post-mortem MRI is valuable as an adjunct to, or instead of, traditional autopsy, providing similar diagnostic accuracy (Thayyil et al., 2013; Kang et al., 2020) and to improve our understanding of congenital neurodevelopmental defects (Vaneckova et al., 2010).

While post-mortem infant MRI overcomes several issues inherent to in vivo MRI scans, it comes with its own unique and complex set of challenges and limitations. The ethical and logistical challenges of subject recruitment results in small sample sizes and the collection of data across restricted age ranges. Furthermore, the acquisition of high resolution post-mortem data is primarily achieved via long scan times. Due to natural tissue degradation during prolonged acquisitions, tissue integrity must be preserved. Most previous post-mortem MRI studies have relied on fixed tissue to tackle this issue (Pfefferbaum et al., 2004; Huang et al., 2009). However, fixation introduces further challenges, such as dramatic reduction in T2 relaxation times and diffusivities (Shepherd et al., 2009; Roebroeck et al., 2019), which negatively impacts signal-to-noise ratio (SNR). This SNR reduction undermines the overall aim of achieving high-resolution imaging.

Ultra-high field imaging of unfixed tissue has significant potential to achieve high resolutions in reasonable scan durations. Several studies have proposed post-mortem MRI of unfixed human fetuses at ultra-high field (>3T) (Sudhin Thayyil et al., 2009; Votino et al., 2012). These studies focused on short (<2 hours) structural MRI scans that aim to provide an alternative to autopsy, demonstrating improved resolution and diagnostic accuracy over conventional MRI. A key challenge for longer scans is the need to minimise degradation of unfixed tissue, requiring that the body be maintained at a low temperature similar to mortuary conditions. Temperature control may be a particular challenge at ultra-high field due to higher radio-frequency energy deposition during imaging.

In this paper, we present the Forget-Me-Not (FMN) study that aims to provide high-resolution dMRI data for connectivity and microstructural estimates in the post-mortem infant. As a substudy of the dHCP, FMN primarily aims to acquire and openly distribute high-quality dMRI data. We use the in vivo dHCP study as a useful comparison throughout, although the aim is for these studies to be both compatible and complementary. For the remainder of the paper, we refer to the previously acquired in vivo dHCP data as the “in vivo” dataset and the novel postmortem dHCP data collected during the FMN study as the “post-mortem” dataset.

We propose an approach that directly addresses the technical challenges involved in using 7T dMRI to image the post-mortem infant brain at high resolution, in situ and unfixed. The acquisition protocol utilizes spin echo excitation and echo-planar readout to enable straightforward comparison with the in vivo dHCP (Hutter et al., 2018) while optimizing acquisition parameters for best image quality based on field strength and the different conditions of in vivo and post-mortem examination. Achieving high resolution imaging data of the intact brain can act as a link beween in vivo dMRI microstructural and connectivity analyses and post-mortem histological findings. The post-mortem diffusion data were acquired using a high b-value multishell protocol to allow the fitting of complex microstructural models, and a relatively large number of diffusion encoding directions to allow high angular resolution tractography. In addition to logistical and technical challenges, the FMN study required careful consideration regarding participant recruitment and research governance; we detail the sensitive recruitment and scanning procedure we developed to enable data to be collected in a single overnight visit shortly after death.

We present results from the first analysable subject recruited to the study. Microstructural properties of the post-mortem cortical plate, subplate, and major white matter tracts were contrasted with previously published findings, both in vivo dMRI and post-mortem histology, to assess the influence of high spatial resolution, low acquisition temperature, and post-mortem tissue degredation effects. Our findings demonstrate that ultra-high field scanning can improve the SNR, and therefore spatial resolution, of neonatal post-mortem imaging. As addtional postmortem dHCP data are collected, it has the potential to facilitate the detailed microstructural assessment of small and intricate brain structures, their developmental trajectories, MRI correlates with histology, and provide a sensitive means to explore the development of both normal and pathological tissue.

## Results

### Participant details and care pathway

Between September 2019 and March 2020, when recruitment was halted for more than a year due to the COVID-19 pandemic, eight infants were identified as eligible for inclusion in the FMN study. The parents of two infants declined to participate, and four eligible infants were not scanned due to technical difficulties (n=2) or practical challenges in coordinating the appropriate research personnel on short timescales (n=2). Two infants were recruited and scanned. Further details regarding the recruitment and transfer to and from the MRI centre are provided in the methods section. Fig. 1 provides a summary of the infants’ care pathway from the neonatal intensive care unit to the MRI centre and to the mortuary. Approximately one week later, parents were sent a letter expressing the FMN team’s appreciation for their participation in the study.

**Figure 1:**
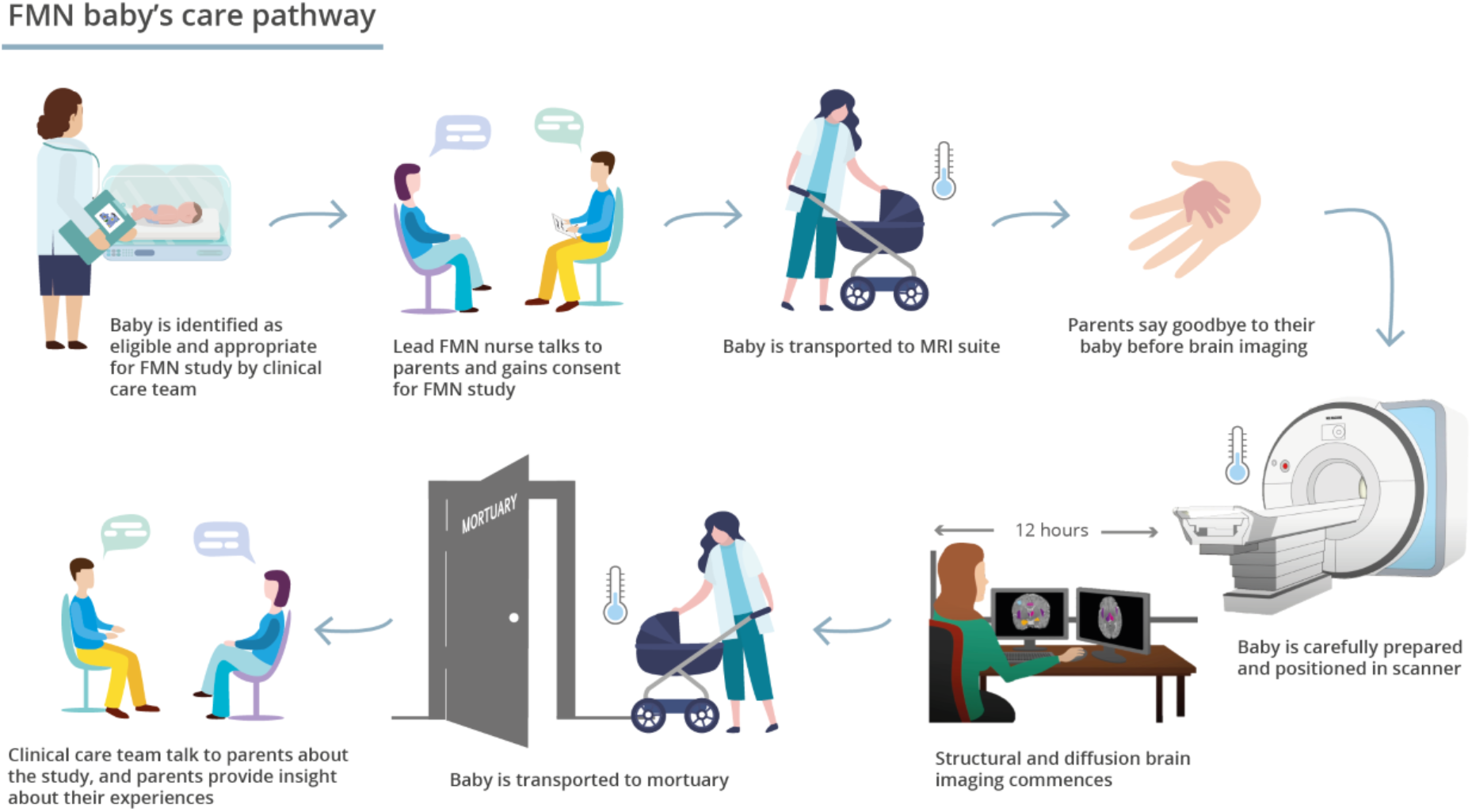
FMN study care pathway. The flow diagram outlines the care pathway for the infants who took part in the study - interaction with parents, transport and cooling system for babies, monitoring during the MRI scan and transfer to mortuary. Parents have an opportunity to see their child again after the child had been transported to the mortuary.

We provide raw data for both infants, but data analysis is only presented in this paper for one infant. The current structural analysis pipeline has not yet been optimised for the other infant, where the significant pathology makes it difficult to use the age-matched brain atlas. The infant presented (male) was born at 23+0 weeks GA, weighing 545g. He had bilateral grade 3 IVH and post-haemorrhagic ventricular dilatation. Following a decision to transition to palliative care, he died on postnatal day 46 at 29+4 weeks PMA and weighing 580g. The infant was scanned within 18h of death.

Parents expressed several reasons for their participation in the FMN study, one of which was the acceptability of the non-invasive nature of the MRI method. Additionally, parents cited altruistic reasons, they commented on their awareness that advances in neonatal care allowing their infants the chance at survival had only come about through research. As such, parents felt it important to contribute to research where able. Along a similar theme, parents were hopeful that their infant’s contribution to research may in some way be useful to the outcome of other infants in future. Thus, taking part provided hope for the future, and parents commented that it had given some meaning to their child’s death.

Study participation did not interfere with the mortuary processes for these infants. Deceased infants often spend periods of time after death on the neonatal unit or in a bereavement suite with their family prior to being taken to the mortuary. As standard of care, cold cots are used in such circumstances to maximise the time that parents can spend time with infants and minimise the impact on any post-mortem assessment. For the purposes of the study, there was a time limit in how long they could spend with the infant prior to the scan taking place. Following the scan, the infants were transferred to the mortuary; however, parents could request that the infant was brought back from the mortuary to the bereavement suite or neonatal unit to allow further time together.

### Data acquisition setup

To facilitate a direct comparison between post-mortem and in vivo data, FMN uses spin echo excitation with EPI readout for dMRI acquisitions, which is consistent with the in vivo dHCP study. Other aspects of the post-mortem study were optimised to provide the best quality data, which resulted in acquisitions that are different to the in vivo dHCP protocols, but is intended to maximise the total information content from the study as a whole. This led to a number of specific decisions regarding acquisition, and a range of technical challenges.

The main challenge for high resolution imaging is obtaining sufficiently high SNR, which scales proportionally to voxel volume. This challenge led to a first key decision for the study, which was to image at higher field strength than the in vivo dHCP study (7T vs 3T). A second major factor dictating SNR in MRI is the radiofrequency (RF) receive coil. In general, image quality improves with coils fit closer to the region of interest, and with increased number of receive channels. Based on the design of the 3T dHCP coil, a 7T RF coil and custom-fitted cradle were designed and built for infant anatomy, as described in the Methods.

The lack of autoregulation in post-mortem imaging increases the risk of tissue heating compared to in vivo scanning. This includes passive tissue heating due to the ambient temperature in the scanner bore rising from heavy use of the gradient coils, and active heating due to RF energy deposition as part of signal formation. Previous post-mortem studies have generally neglected heating, and either used fixed tissue that is less susceptible to damage, or scanned unfixed tissue only for a short time such that temperature rises are likely to be modest. However, active heating is a particular concern for FMN, because (a) the study is conducted at 7T, and RF energy deposition increases quadratically with field strength, (b) several sequences used in this study are generally of high energy deposition (in particular, dMRI using simultaneous multi-slice spin-echo imaging), and (c) the long scan duration used in this study leads to more heat accumulation. Due to the sensitive nature of the study, we had to guard conservatively against any tissue damage due to heating. A secondary concern is that tissue heating alters water diffusivites during the scan, requiring modelling to remove the confounding effects of time-varying water diffusivity. We conducted an investigative study of an unfixed porcine brain, finding tissue temperature increases of 5 °C during a 6-hour diffusion scan. We therefore developed an active cooling system with the aim of stabilising the infant’s body temperature to a value between 5 and 10°C, which is consistent with mortuary conditions, and accept no more than 1-2 °C variation during a given scanning modality. System specifications, design and evaluation are described in Supplementary Information S1.

Compared to in vivo imaging, post-mortem dMRI provides both unique opportunities (e.g. much longer scan times) as well as challenges (e.g. changes to tissue properties). Many post-mortem dMRI studies leveraged the former to address the latter by employing diffusion-sensitising pulse sequences that are not feasible in vivo (Fritz et al., 2019; McNab et al., 2009; Miller et al., 2012). However, these approaches would be less straightforwardly compatible with the in vivo dHCP study, which acquired conventional single-shot diffusion-weighted spin-echo data. We therefore opted to use a diffusion-weighted spin-echo sequence with a modified imaging readout: specifically, a readout-segmented acquisition (Porter and Heidemann, 2009). This method achieves high-resolution images without incurring problematic levels of distortion. While this approach decouples image distortion from resolution in a way that single-shot methods cannot, it also incurs a cost of increased scan time. We mitigated this challenge using simultaneous multi-slice acquisition (Larkman et al., 2001; Setsompop et al., 2013; Frost et al., 2015). This relatively new acquisition method was not yet available on our vendor-provided platform, requiring custom implementation including optimised radiofrequency pulses, automated raw data transfer and offline image reconstruction.

dMRI protocol optimisation had to be undertaken in light of two conflicting considerations. First, to our knowledge the specific scan conditions of the FMN study had not been investigated before, resulting in a lack of information about T1, T2 and mean diffusivity for unfixed post-mortem infant brain at 7T. Knowledge of these properties is critical for protocol optimisation. Second, the number of infants recruited was expected to be low, making each recruitment extremely valuable and raising concerns about the appropriateness of using recruited babies for detailed piloting for protocol optimisation. We therefore adopted a strategy to optimise acquisition protocols based on predicted tissue parameters extrapolated from literature, while retaining the possibility to adjust some critical parameters (e.g., b values) based on a short pilot at the beginning of the first infant scan. This approach aimed to maximise the likelihood of obtaining usable data in the first infants whilst taking into consideration the high degree of uncertainty regarding tissue parameter predictions.

The imaging protocol used to acquire the first two infants’ data includes T1- and T2-weighted and dMRI protocols. Here, we present the T2w and dMRI data. T2-weighted structural data were acquired at 0.4 mm isotropic resolution (compared to 0.8 mm resolution for in vivo dHCP data). dMRI data were acquired at 0.8mm isotropic resolution (compared to 1.5mm resolution for in vivo dHCP data). Similarly to the in vivo dHCP data, we acquired dMRI data with three shells but with evenly spaced b-values adjusted to account for reduced diffusivities: b=3000, 6000 and 9000 s/mm^2^ (compared to b=400, 1000 and 2600 s/mm^2^ for in vivo dHCP). The protocol was optimised for data quality achievable in an overnight scan session, which is described in Supplementary Information S2.

### Structural and diffusion data quality assessment

Cranial ultrasound images of the infant taken prior to death reveal evidence of bilateral IVH grade 3, post-haemorrhagic ventricular dilatation, and parenchymal extension in the left hemisphere (Fig. 3 Ultrasound). These pathological details are visible with finer detail and improved contrast on the post-mortem structural T2-weighted images at 7T (Fig. 3 Post-mortem T2w). The gain in structural detail in the post-mortem structural images compared to in vivo is highlighted by comparison to an age-matched subject from the in vivo dHCP dataset (Fig. 3 In vivo T2w). This reflects the higher spatial resolution (0.4 mm post-mortem vs 0.8 mm in vivo), achievable due to the higher field strength and longer scan time, and lack of motion-related artefacts. The post-mortem T2w image provides clearer delineation of boundaries between different tissue structures, for example, the deep gray matter structures forming the thalamus and basal ganglia.

**Figure 2:**
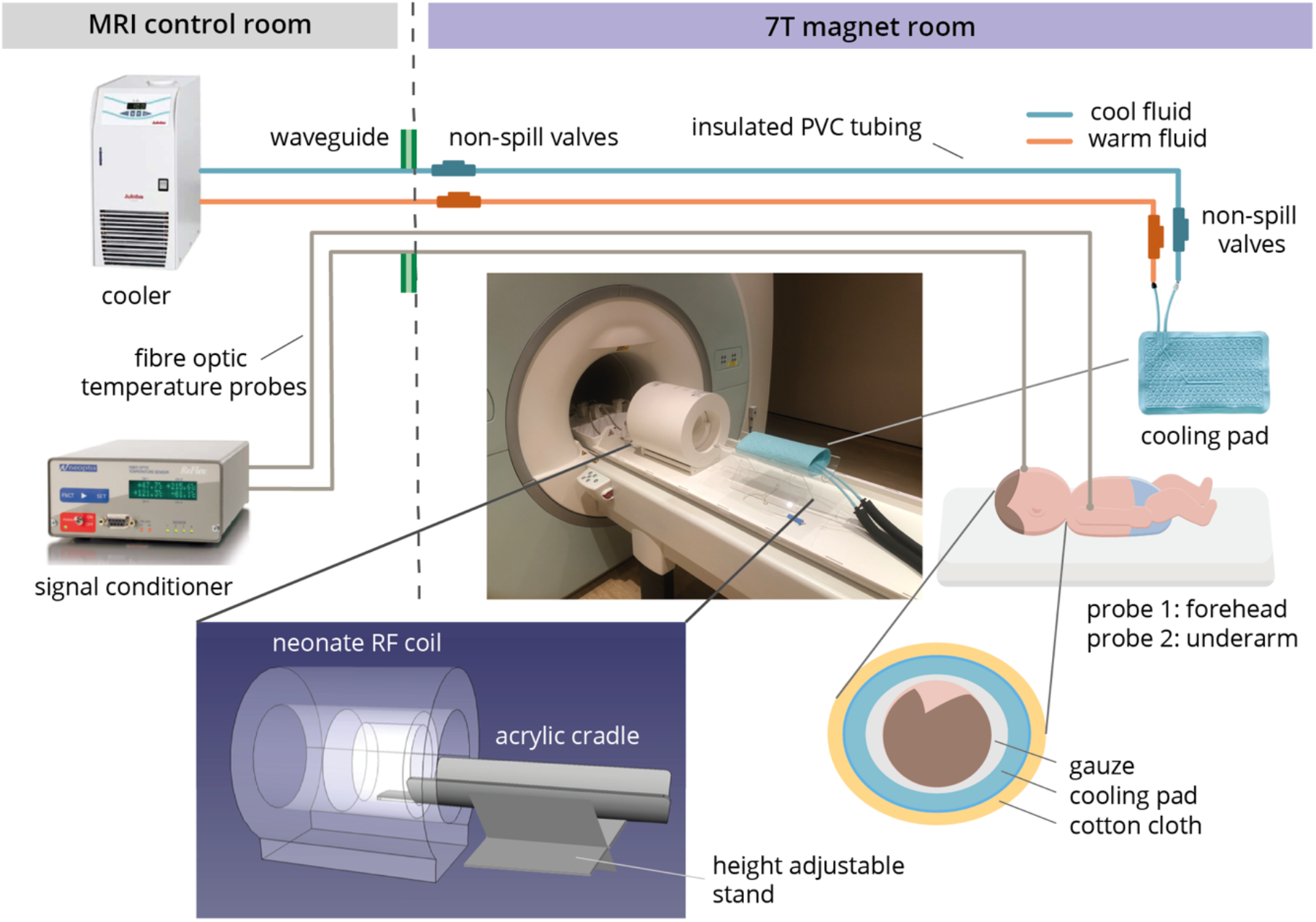
Data acquisition setup. Post-mortem neonate data are acquired at 7T with a RF coil and custom-fitted cradle specially designed for infant anatomy. An active cooling system was built to stabilize infant body temperature to a value between 5°C and 10°C during a given scan modality. Infant body temperature is monitored in real time during the scan using two fibre optic temperature probes attached to the infant’s forehead and underarm, respectively.

**Figure 3:**
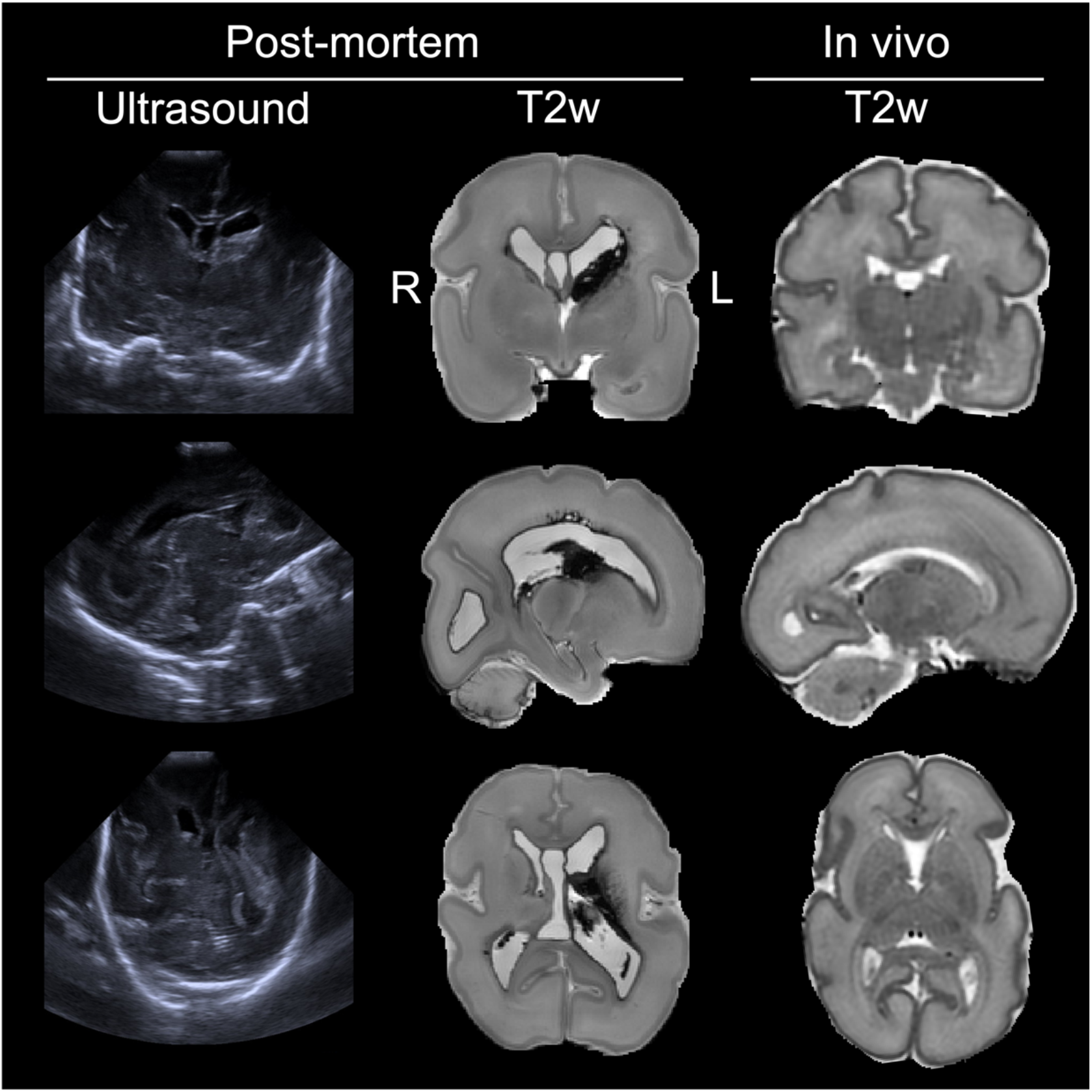
Structural data quality assessment. For the post-mortem dHCP subject, the ultrasound images were acquired prior to death, while the T2w structural MRI is post-mortem. The in vivo T2w data are from an age-matched infant from the in vivo dHCP database. The post-mortem and in vivo T2w images were acquired at 0.4×0.4×0.4mm^3^and 0.8×0.8×0.8mm^3^ spatial resolutions, respectively. Note the in vivo T2w images were interpolated to 0.5×0.5×0.5mm^3^ in post-processing. The post-mortem images were acquired using 3D T2w-SPACE sequence and the in vivo images were acquired using 2D multi-slice FSE sequence with motion corrected reconstruction.

Fig. 4 compares the dMRI data between the post-mortem subject and the exemplar age-matched in vivo subject. Despite the 6.6-fold smaller voxel volume of post-mortem images compared to in vivo (0.512 vs 3.375 mm^3^), the post-mortem data have sufficient SNR to support robust diffusion model fitting. This is true for both the representational diffusion kurtosis imaging (DKI) model (Jensen et al., 2005) and the biophysical neurite orientation dispersion and density imaging (NODDI) model (Zhang et al., 2012), although caveats regarding the NODDI parameter estimates are discussed later. The quality of model fits is clear from the relatively high consistency in overall spatial patterns of model parameters between the post-mortem and in vivo data. Nevertheless, there are notable differences. Our post-mortem data exhibited higher mean kurtosis and higher intra-neurite volume fraction than the in vivo data (Fig. 4b,c; note the differences in colourbar scale). Additionally, the post-mortem mean diffusivity is considerably lower than that of the in vivo data, consistent with many previous post-mortem studies in both fixed and unfixed tissue. The mean diffusivity of CSF in the post-mortem data is 2.8 times lower than that in the in vivo data, which is consistent with the lower temperature of the post-mortem scan (10°C rather than 37°C, leading to a ~2X reduction of water diffusivity). Notably, a much larger difference of mean diffusivities is observed in brain tissue (~10 fold lower in the post-mortem than in vivo). These differences might be explained by death related changes such as postmortem cell swelling (Toorn et al., 1996; Xiao et al., 2020).

**Figure 4:**
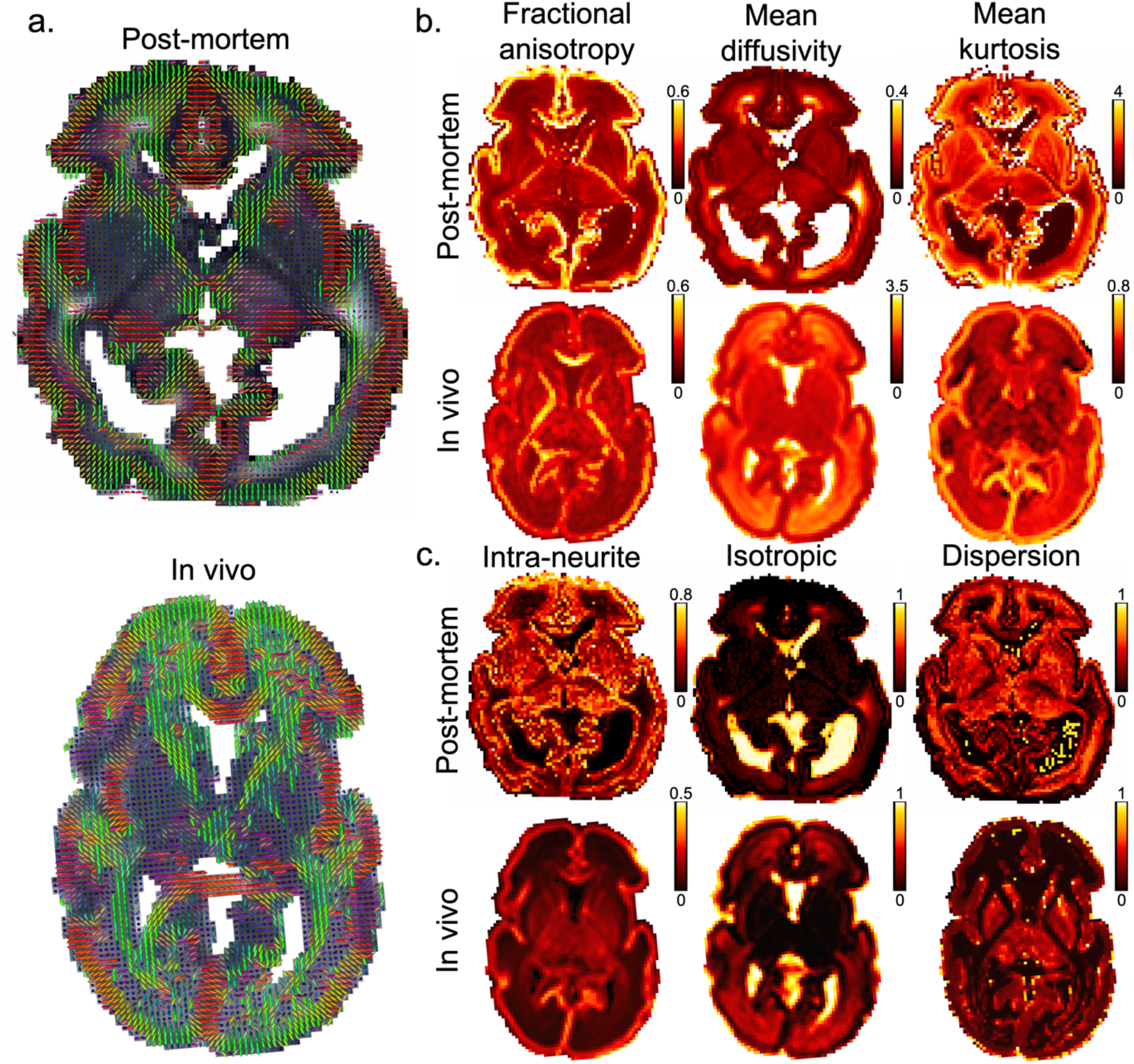
Microstructural data quality assessment. In all panels, there is a high degree of qualitative correspondence in spatial patterns between the post-mortem data and the exemplar age-matched in vivo subject. In general, greater microstructural detail can be resolved in the post-mortem data relative to the in vivo data due to the higher spatial resolution. Note the acquisition time for the post-mortem data is much longer than the in vivo data (7h v.s. 19min). a. Principal diffusion direction from DKI model fit. b. Mean diffusivity (μm^2^/ms), fractional anisotropy, and mean kurtosis parameters from the DKI model fit. c. Intra-neurite volume fraction, isotropic volume fraction, and orientation dispersion index parameters from NODDI model fit. Quantitatively, overall spatial patterns of DKI and NODDI parameters are consistent between subjects. However, in the post-mortem data, the mean diffusivity values are noticeably lower, while the mean kurtosis and the intra-neurite volume fraction values are noticeably higher, than the in vivo data. Note the change of scale bar between the post-mortem and in vivo maps. The post-mortem data has a resolution of 0.8mm isotropic, and the in vivo data were interpolated to 1.17×1.17×1.5mm^3^ in post-processing.

In the results that follow, we analyse the microstructural properties of the cortical plate, subplate, and white matter tracts. An over-arching goal is to assess the value that the post-mortem dHCP data adds to visualising and analysing infant brain microstructure, the suitability of this data to act as a bridge between histological findings and the in vivo dHCP dMRI data and the potential to use the post-mortem data to investigate pathology and death related tissue changes. To facilitate interpretability and cross-dataset comparisons, we restricted analyses to the right hemisphere only, to mitigate the influence of significant overt pathology in the left hemisphere of the post-mortem subject (Fig. 3).

### Cortical plate radial microstructure

Cerebral cortex development takes place in an ordered manner that involves distinct cellular structure changes at different gestational times. Early in development, the orientation of glial fibers and the apical dendrites of pyramidal cells are primarily radial with respect to the cortical surface. Diffusion MRI studies of non-human primate (Kroenke et al., 2007, 2005) and human brains (Maas et al., 2004; McKinstry, 2002) have found a predominantly radial principal diffusion direction in cortex with high anisotropy at early development that reduces with cortical maturation.

The post-mortem dHCP data show radial microstructural organisation of the cortical plate (fig. 4a), the precursor to developed cortex, which is consistent with histological studies (Marín-Padilla, 1992) and dMRI (McKinstry, 2002) conducted at this gestational age. This cortical radiality can be quantified with the radiality index (McNab et al., 2013) (i.e., dot product of the principal eigenvector of the diffusion tensor and the surface normal, Fig. 5b). In the post-mortem imaged brain, most of the cortex has a radiality≈1, but the radiality is consistently reduced at the gyral walls. This is likely due to partial volume effects, where fibres that turn sharply to enter the cortical plate cannot be resolved even at this high resolution because of the thin cortical plate in neonates.

**Figure 5:**
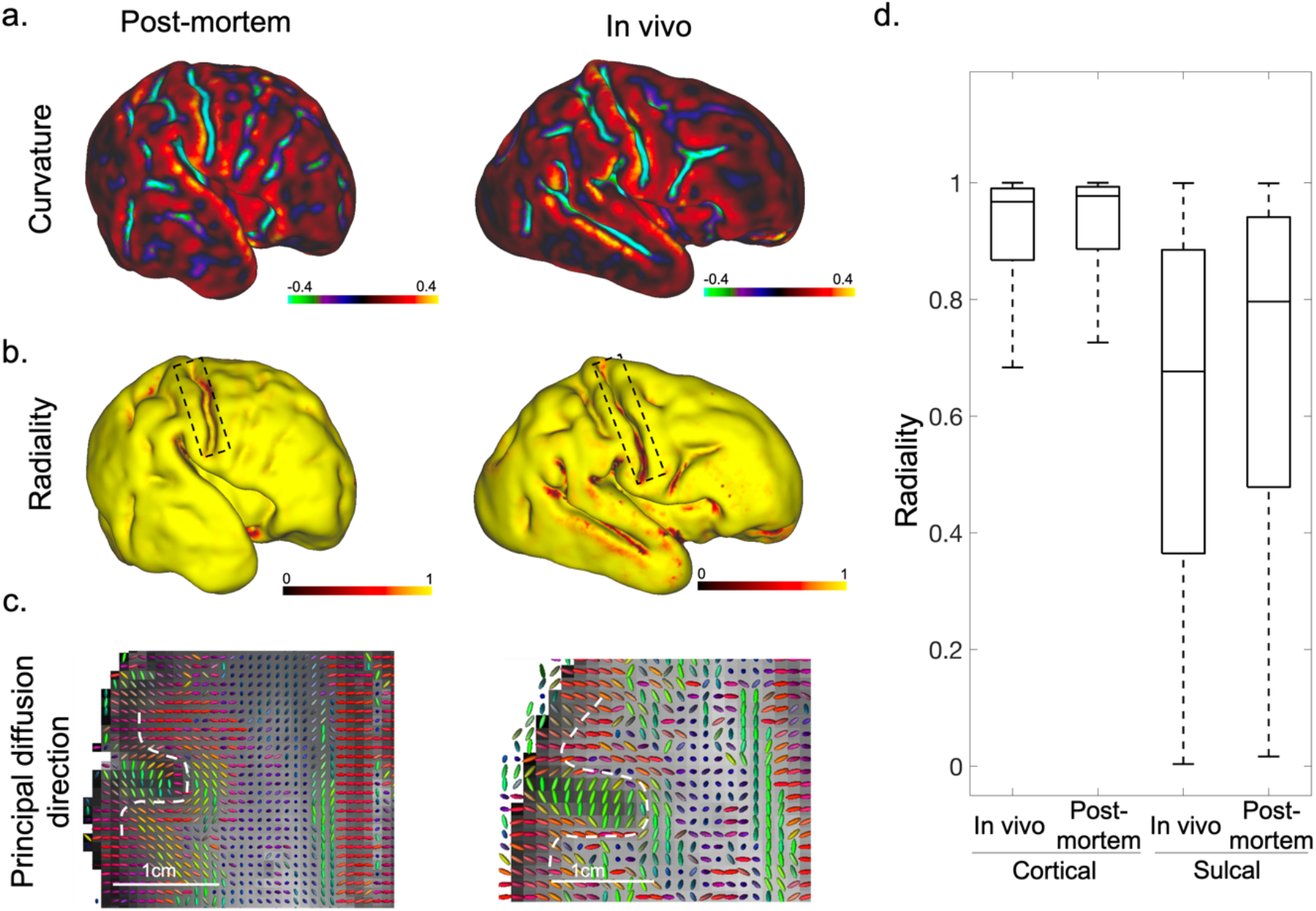
Cortical plate microstructure. (a). Cortical curvature measured at mid-thickness of cortical ribbon from the post-mortem subject and an age-matched in vivo subject. (b). Cortical radiality (dot product of the principal eigenvector of the diffusion tensor and the surface normal) calculated at mid-thickness of cortical ribbon from the same subjects. Overall, both in vivo and post-mortem brains exhibit high radiality values across the cortex. In regions of high curvature, such as at the fundus of the central sulcus (black box), the post-mortem data have higher radiality than the in vivo data. (c). Principal diffusion directions for an axial plane containing central sulcus, the dashed white line indicates the gray-white matter boundary. The principal diffusion directions at the fundus are radial in the post-mortem data but tangential in the in vivo data. Note the in vivo data were upsampled from 1.5×1.5×1.5mm^3^ to 1.17×1.17×1.5mm^3^ in post-processing. (d). Box plots of cortical radiality and sulcal radiality calculated from data points at the mid-thickness of cortical ribbon from the post-mortem subject and five age-matched in vivo subjects. Both the post-mortem and in vivo data exhibit a high overall radiality across the cortex with similar drops in radiality in sulcal regions. The post-mortem data exibit slightly higher radiality values in sulci, likely reflecting the benefits of higher spatial resolution.

The radial microstructural organisation across the cortex of the post-mortem data is strikingly similar to cortical radiality of in vivo subject’s data (Fig 4a). Compared to the in vivo subject, the post-mortem subject’s data have quantitatively higher radiality in several regions, such as at the fundus of the central sulcus (Fig. 5b black boxes and 5c) as well as several other regions of high curvature (sulcul fundi and, to a lesser extent, gyral crowns), which can be seen by comparing Figs. 5a and 5b. The higher radiality in high curvature regions in the post-mortem data likely reflects the benefits of increased spatial resolution achieved with longer scan time.

Using the group average of in vivo data from 5 subjects (for details see Supplementary Table S2), these similarities in cortical plate radiality patterns were also visible between the post-mortem and in vivo data. Fig. 5d compares radiality across cortex versus sulcal fundi in the post-mortem and in vivo data. Fundi masks were generated by thresholding the curvature map to isolate surface regions with curvature value less than −0.4. In the in vivo data, there was a 30% difference between the median cortical radiality (0.967) and the median sulcal radiality (0.677); while in the post-mortem data, there was a comparable 20% difference between the median cortical radiality (0.977) and the median sulcal radiality (0.796). Again the higher radiality values in the post-mortem data are likely a reflection of the benefits of higher spatial resolution.

### Subplate microstructure

We characterised variability in diffusion properties across the subplate, a prominent layer immediately adjacent and deep to the cortical plate. This developmentally-important transient layer acts as a staging area for neuronal fibres, is crucial for establishing cortico-cortical and cortical-thalamic connections, and is characterised as having low cell density and high synapse density and extracellular matrix content (Kostović et al., 2019). To investigate the microstructure properties in subplate, we assessed DKI parameter intensity profiles as a function of tissue depth that extended 4mm inward from the cortical plate-subplate interface (Fig. 6). The median parameter values across the cortical plate are plotted at 0mm.

**Figure 6:**
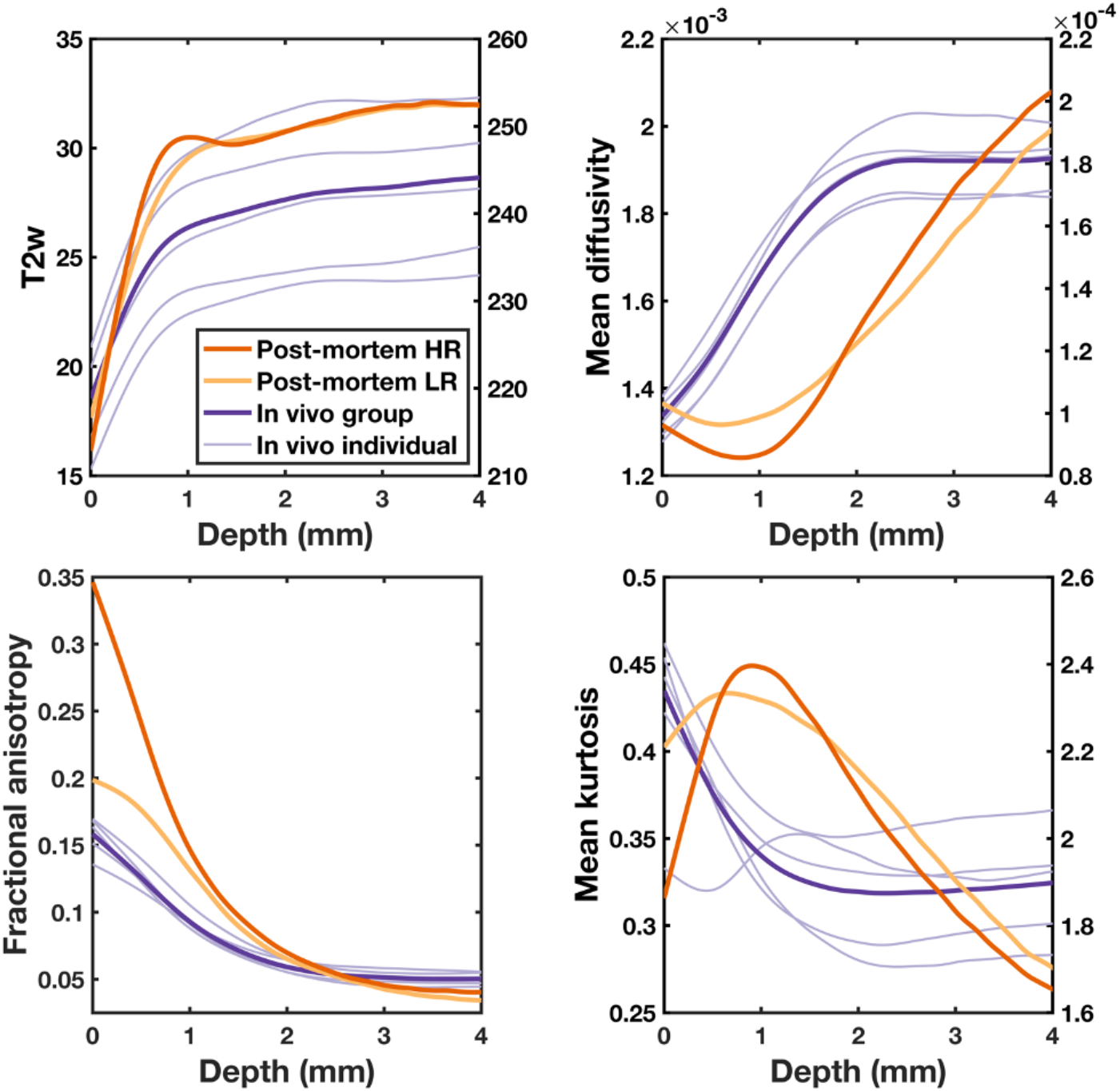
Subplate microstructure. T2w image intensity and DKI parameters plotted as a function of the distance from the cortical plate for (i) high resolution (0.8mm/0.4mm isotropic for dMRI/T2w images) post-mortem data (red), (ii) low resolution (1.5mm/0.8mm isotropic for dMRI/T2w images) post-mortem data (yellow) generated by downsampling the high-resolution post-mortem data, (iii) five age-matched subjects scanned in vivo (light purple), (iv) group median of the five subjects scanned in vivo (purple). The median parameter value is taken for a given depth, and a smoothing spline curve was fit to the data. The group median of the five subjects scanned in vivo was calculated as the median parameter value across all subjects for a given depth. The resolution of in vivo dMRI and T2w images are 1.5mm and 0.8mm isotropic, respectively. For clear visulisation, T2w, mean diffusivity, and mean kurtosis values are plotted using separate scales per dataset, with in vivo data scaled according to the left y-axis and post-mortem data scaled according to the right y-axis. Abbreviations: Post-mortem HR = Post-mortem high resolution; Post-mortem LR = Post-mortem low resolution; In vivo group = group median of in vivo dHCP data, In vivo individual = individual in vivo dHCP subject data. The mean diffusivity is expressed in mm^2^/s.

The T2w image intensity was used to localise the superficial subplate as a hyperintensity peaking approximately 1mm from the interface with the cortical plate, in line with previous histologically-validated findings (Pogledic et al., 2020). This hyperintensity peak in T2w images was not visible in the in vivo data or the post-mortem data after downsampling to the spatial resolution of the in vivo data. The FA (fractional anisotropy) profiles for both post-mortem and in vivo data have highest FA in the cortical plate, decreasing in magnitude with depth. The high resolution post-mortem data demonstrated the greatest difference in FA between cortical plate and deeper layers, with the downsampled post-mortem profile more closely matching the in vivo profile. Taken together, it is clear that high spatial resolution is helpful in identifying the superficial subplate from T2w data and to accurately characterise the high FA cortical plate from the much lower FA subplate.

In the post-mortem dMRI data, mean diffusivity and mean kurtosis values demonstrate peaks or troughs at a depth of approximately 1mm, corresponding to the superficial subplate identified by the hyperintensity peak in the T2w image. Mean diffusivity reaches its minimum and mean kurtosis its maximum in the region of the T2w peak, suggesting higher complexity of microenvironment in the subplate. The subplate parameter peaks and troughs in the post-mortem profiles are not visible in the in vivo data. This likely reflects pathology and tissue changes due the death, as downsampling the post-mortem data to the lower resolution of in vivo data did not eliminate these features.

### White matter microstructure

Finally, we examine the microstructural properties of white matter tracts. In neuroimaging tractography studies, it is typical to classify tracts into functional categories based on the nature of the tracts’ origin and termination points. Three functional categories were of relevance to this study: projection tracts connecting cortex–spinal cord, cortex-brainstem, and cortex-thalamus; association tracts connecting cortex-cortex within a hemisphere; and limbic tracts connecting limbic system structures (Bastiani et al., 2019; Ouyang et al., 2019). Tracts are estimated in the right cerebral hemisphere using probabilistic tractography and the protocols developed as part of the dHCP (Bastiani et al., 2019). The 11 tracts included in our analysis included five projection tracts – corticospinal tract, auditory radiations, and anterior, posterior, and superior thalamic radiations; three limbic tracts – fornix, cingulate and hippocampal cingulum bundle; and three association tracts – uncinate, superior longitudinal, and inferior longitudinal fasciculus (Fig. 7a).

**Figure 7:**
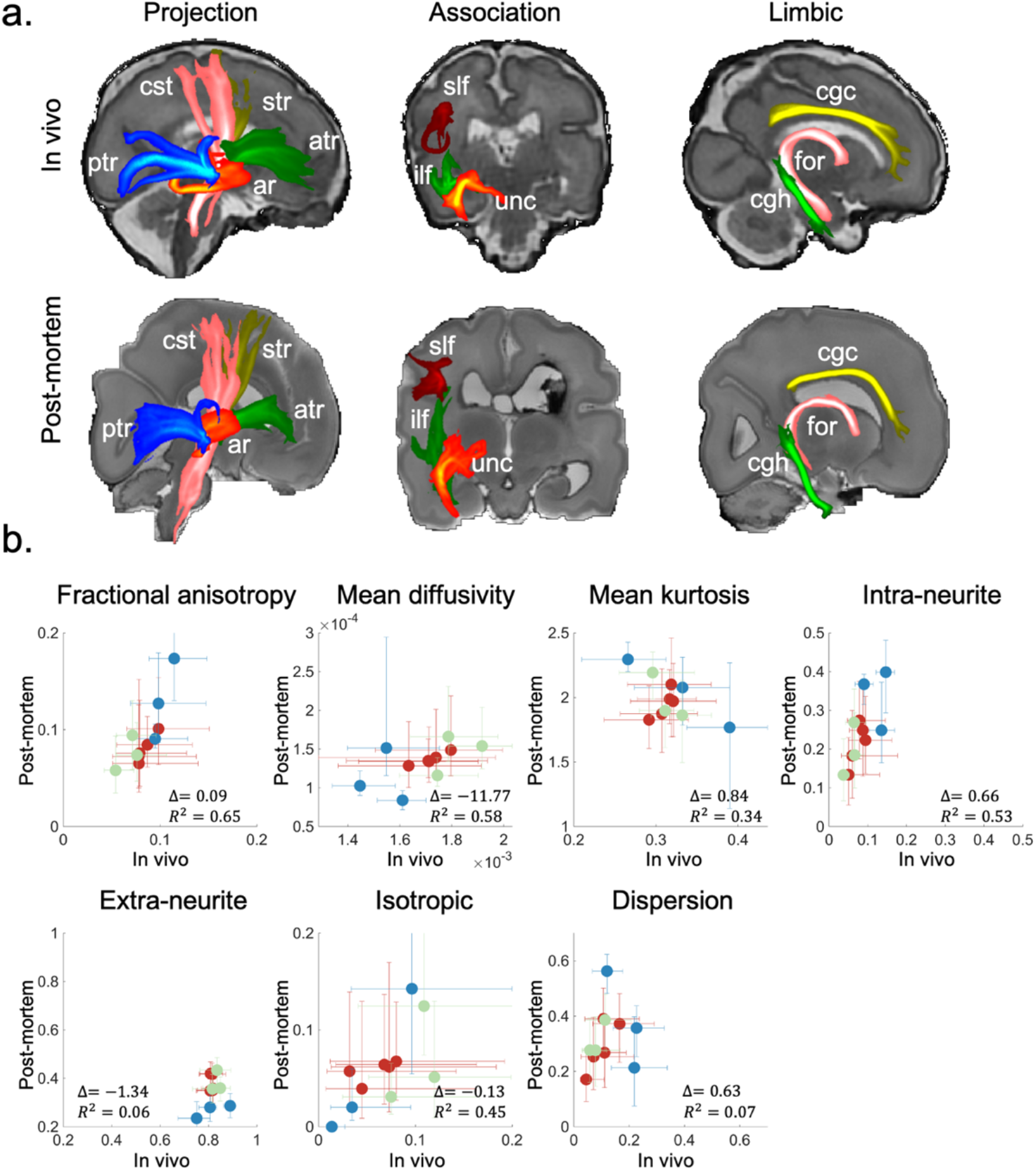
White matter microstructures. (a). Maximum intensity projections of 11 tracts from the in vivo and post-mortem data. All tracts are visualised using the same thresholds (0.005, 0.05). (b). The median DKI and NODDI parameters were calculated per tract for three categories of white matter pathways: association (green), limbic (blue) and projection (red). Scatter plots display diffusion parameters from post-mortem v.s. in vivo dHCP data. Δ:relative difference between tract-averaged parameter values in post-mortem and in vivo dHCP data ((post-mortem - in vivo)/post-mortem), indicating global difference in parameter scaling. R^2^: coefficient of determination for linear regression (equivalent to squared correlation coefficient), indicating cross-dateset similarity in tract-to-tract spatial variability patterns. Error bars indicate interquartile range (i.e. difference between 75^th^ and 25^th^ percentiles). The mean diffusivity is expressed in mm^2^/s. Abbreviations: ar = acoustic radiation; atr = anterior thalamic radiation; cgc = cingulate gyrus part of the cingulum; cgh = parahippocampal part of the cingulum; cst = corticospinal tract; for = fornix; ilf = inferior longitudinal fasciculus; ptr = posterior thalamic radiation; slf = superior longitudinal fasciculus; str = superior thalamic radiation; unc = uncinate fasciculus.

Fig. 7b summarises DKI and NODDI parameter estimates for white matter tracts. Comparing spatial patterns across tracts, the post-mortem data and in vivo data have similar tract-to-tract variability for fractional anisotropy, mean diffusivity and intra-neurite fraction with coefficients of determination R^2^>0.5. Comparing global parameter scaling between datasets, the NODDI results suggest that compartment fractions are globally inverted between the datasets, with the post-mortem data having consistently larger intra-neurite (Δ((post-mortem - in vivo)/post-mortem)=0.66) and lower extra-neurite volume fractions (Δ=−1.34) compared to the in vivo data. This would be consistent with cellular swelling (cytotoxic edema) reducing the extracellular space. Orientation dispersion parameters are generally higher in the post-mortem data than the in vivo data, which may also reflect death related tissue changes.

Caution also needs to be taken when interpretating these results due to the established deficiencies in the NODDI model (Jelescu et al., 2016). In particular, we found the dispersion parameter is highly variable depending on the diffusivity value used in the fitting, which we needed to predetermine but have no accurate knowledge of. The inaccuracies in diffusivity can lead to increased/decreased dispersion estimation in NODDI fitting (Yi et al., 2019). Notably, while we found dispersion estimates to depend strongly on the assumptions of the NODDI model, the overall patterns of intra- vs extra-neurite fractions was remarkably invariant to these assumptions, which could be a robust metric for investigating tissue changes due to death or pathology.

As a final exploratory analysis, we assessed the ability to segregate these three tract categories (projection, association and limbic) based on the median parameter value per tract for DKI and NODDI parameters. Scatter plots of pairs of DKI parameters result in clusters of tracts that can be broadly visualised from both the post-mortem data (Fig. 8a) and the in vivo group data (Fig. 8b), suggesting that tracts do exhibit some segregation based on function. To explore this clustering more quantitatively, we utilize linear discriminant analysis (LDA) for different sets of diffusion measures (DKI alone, NODDI alone, and both DKI and NODDI). For a given set of features, we use LDA to calculate a single discriminant (multi-parameter combination) that provides maximal separability between tract categories. We can then quantify the degree of clustering using the linear discriminant ratio (LDR, or Fisher’s linear discrimant: the ratio of inter- to intra-class variances), which is the same quantity that is maximised in LDA.

**Figure 8:**
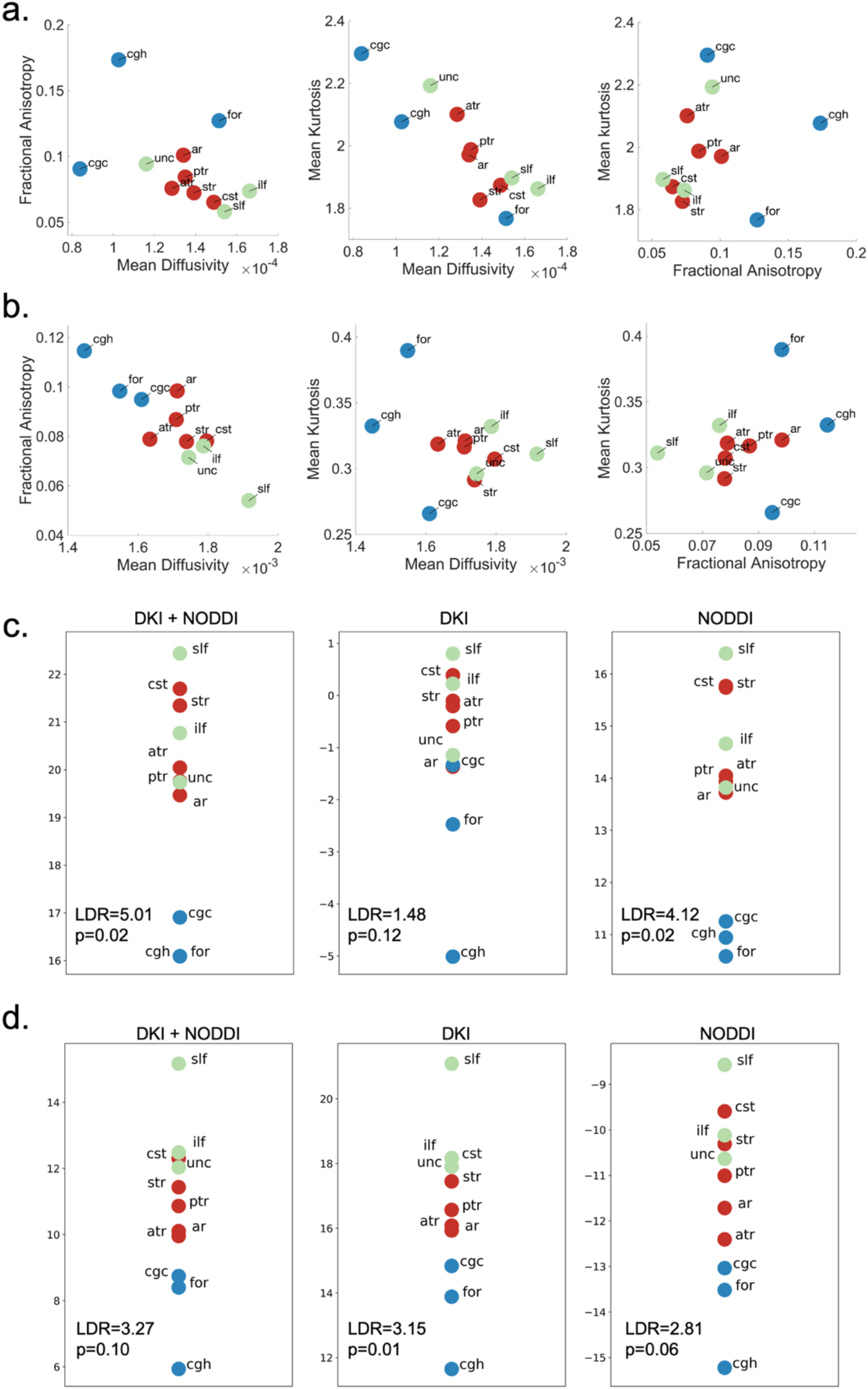
White matter tract segregation. Scatter plots of pairs of DKI parameters (Mean diffusivity v.s. Fractional anisotropy, Mean diffusivity v.s. Mean kurtosis and Fractional anisotropy v.s. Mean kurtosis) from the post-mortem data (a) and in vivo group data (b). Tract segregation of post-mortem data (c) and in vivo group data (d) projected to the optimal one-dimensional linear discriminant feature space. The dots represent median DKI and NODDI parameters calculated for three categories of white matter pathways: association (green), limbic (blue) and projection (red). P values are calculated from permutation analyses. The mean diffusivity is expressed in mm^2^/s. Linear discriminant ratio (LDR) is the ratio between inter- and intra-class variances.

Results from the post-mortem data (Fig. 8c) and the in vivo group data (Fig. 8d) show that a single LDA derived feature vector (discriminant) is able to achieve clear separation between limbic and non-limbic tracts, but less able to separate projection and association tracts. The consistency between the post-mortem and the in vivo data suggests the pathology and death related tissue changes are sufficiently consistent as to maintain the relationship between tracts. We used permutation testing to calculate p-values, showing high confidence over chance on the results for most, but not all, evaluations. Permutation testing is particularly helpful in comparing LDR values across the different parameter sets, where discriminants trained with a larger number of input features will be more prone to overfitting. In these cases, random permutations will also be overfit and achieve high LDR values, which then results in higher p-values. For example, in the dHCP data, the DKI (3-parameter) and DKI+NODDI (7-parameter) models have similar LDR values (3.15 and 3.27, respectively) but quite different p-values (0.01 and 0.12, respectively). It is important to note that even when comparing post-mortem and in vivo clustering derived from the same set of diffusion metrics (e.g. DKI parameters) the actual discriminants (LDA feature vectors) are derived from the two datasets separately and are unlikely to reflect the same information.

## Discussion

The post-mortem Forget-Me-Not dHCP study aims to acquire and openly distribute high-quality diffusion imaging of unfixed post-mortem infant brains that will provide insight into developing brain connectivity and microstructure. Furthermore, the data are designed to enable direct comparison with the in vivo dHCP dataset. The post-mortem data appeared to resolve fine brain structures and to detect cortical radiality in regions with high curvature, results that suggest a direct benefit of the increased spatial resolution. While the post-mortem data have greatly improved spatial resolution relative to the in vivo data, the post-mortem brain features macroscopically visible pathologies, and our dMRI analysis results suggest that there are further microstructural changes, such as cell swelling and autolysis, which are expected consequences of pathological and death-related processes. These effects introduce challenges in translating microstructural findings from post-mortem to in vivo diffusion imaging data, which the FMN study aims to address as further data are collected.

One major component of the FMN study is the care pathway, which was designed around unique challenges associated with conducting a post-mortem infant imaging research study. The death of an infant represents a profound tragedy and highly stressful time for the families involved, and it is vital that there is careful ethical consideration of recruitment of parents for participation in a research study. For this particular study, recruitment was initiated by the infant’s clinical team, independent of the study, who identified infants eligible and appropriate for recruitment. The lead FMN research nurse (F.A.), who gave a verbal explanation of the purpose and goal of the study and recruited the family into the study, had undergone appropriate bereavement training. Parents that participated in the FMN study cited reasons for participation including: appreciation that previous research formed the foundation of the clinical care they and their infant received, the hope that data collected might help future families, and a sense of meaning to their infant’s death that they felt due to participating. These are similar to the expressed views of parents in previous post-mortem research studies in the UK (S. Thayyil et al., 2009). Previous studies have indicated that participation in research is not usually harmful and often positively received by bereaved families (Butler et al., 2018).

Other aspects of the FMN study that had positive effects on recruitment to date were the non-invasive nature of MRI imaging and the in situ scanning at low temperatures similar to mortuary conditions, but unfixed. These observations are consistent with findings suggesting that fewer than 50-60% of parents consent to invasive autopsy, mainly owing to concerns about body disfigurement (Kang et al., 2020). However, without fixation, the scanned tissue is in a more volatile state. In the future, scanning the brain pre- and post-fixation and acquiring subsequent histological data would provide valuable insight to link histology with dMRI assessment of microstructure.

A second major thread of work in the FMN study is the development of the experimental setup and imaging protocols for high resolution dMRI of the unfixed post-mortem neonate brain. The constraints of the FMN study led to a number of technical challenges that required different approaches to our previous post-mortem imaging in adult brains. A first challenge was to achieve higher spatial resolution whilst minimising EPI-related distortion and blurring. The most common approach to this problem is to use single-shot EPI with in-plane acceleration; however, we found our coil to be unable to deliver the acceleration necessary to meet our target resolution of 0.8mm. Instead, we utilised readout-segmented EPI, which reduces distortion by breaking up the 3D volume into readout segments (Porter and Heidemann, 2009). While each individual segment is shorter, the entire 3D volume takes longer to acquire with readout segmentation. In order to acquire our target number of diffusion volumes in an overnight scan, we needed to then reduce the scan time per volume. This was achieved by integrating more modest accelerations in both the slice direction (simultaneous multi-slice imaging with MB=2) and in plane (phase-encoding parallel imaging with R=2). By integrating these three imaging techniques (readout segmentation, multi-slice acceleration and in-plane acceleration) (Frost et al., 2015), along with the use of 7T and a bespoke 16-channel neonatal head coil, we were able to meet the acquisition target and provide high quality dMRI data.

A concern that was relatively unexplored at the start of our study, but which proved to be a major technical challenge, was tissue heating. Previous work in post-mortem dMRI of fixed tissue has established that long scan times coupled with the need for high b-value lead to significant gradient heating, which raises the ambient room temperature and can passively heat samples (Miller et al., 2011). Our study conditions introduced additional concerns regarding RF energy deposition that is particularly problematic at ultra-high field (since energy levels scale with the square of field strength) and in post-mortem tissue (where there is no auto-regulation to mitigate heating and unfixed tissue is particularly susceptible to damage). Tissue damage from heating would be problematic both ethically and with respect to later autopsy. To overcome this challenge, we developed an active cooling system, which enables stable and low body temperature during scanning of dMRI data and is easy to operate. A secondary benefit of stable body temperature is that tissue properties such as T2 and ADC will also remain stable during the scan. The design of this cooling system may be a useful resource for similar studies in the future.

Although the main focus of the FMN study is to estimate diffusion parameters, other tissue parameters such as relaxation times can offer important but complementary information that can be useful in providing a comprehensive depiction of tissue structures. In the first two FMN scans, we piloted quantitative multi-parameter mapping sequences (Weiskopf et al., 2013). While this data was not usable in the first babies due to image artefacts, these will be straightforward to address and we aim to include this protocol in future scans. We thus anticipate future datasets providing T1, T2*, proton density and magnetic transfer saturation at 0.8mm isotropic resolution in a ~1.5 hour scan.

A major aim in many post-mortem MRI studies is the direct comparison with histological data, enabling validation of imaging results. To date, FMN has not attempted to recruit brain donation, which would enable histology data to be collected, but might affect parents’ willingness to participate. While we feel it is unlikely that many parents would consent to brain donation, we will aim to explore this possibility in the future. Acquisition of histological data along with imaging data in a subset of future recruits may be possible, and would greatly improve the value of the resource.

Overall, the structural and microstructural maps for the in vivo and post-mortem dHCP data were qualitatively similar, with the post-mortem data exhibiting the desired improvements in spatial resolution (Figs. 3–4). The radial organisation of the cortical plate around 29 weeks GA is visible in histological preparations (Marín-Padilla, 1992) and results in the cortical plate exhibiting greater radial organisation of principal diffusion directions (McKinstry et al., 2002) and higher FA values (Ball et al., 2013) than the deeper subplate layer in dMRI images. Additionally, the increased extracellular matrix content of the subplate relative to the cortical plate is visible in histological preparations (Kostović et al., 2019; Pogledic et al., 2020) with the superficial subplate corresponding to a hyperintense layer in T2w images (Pogledic et al., 2020). Using these histological features as “gold standard” benchmarks, the post-mortem data demonstrate the capability to faithfully reflect the underlying histology thanks to the high spatial resolution.

High cortical plate radiality and FA values are seen in both the post-mortem and in vivo dHCP data. However, the post-mortem data exhibit more homogeneous cortical radiality (Fig. 5) and greater FA contrast between the cortical plate and deeper structures (Fig. 6). Additionally, the superficial subplate T2 hyperintensity was only seen in the post-mortem data (Fig. 6). Downsampling the post-mortem data to allow comparison with the in vivo data without a change in resolution, the T2w hyperintensity disappears and the FA contrast between cortical plate and subplate dramatically reduces. These results demonstrate the advantage of high resolution data despite long scan hours, which enables more detailed depiction of small structures. The patterns in the downsampled postmortem data were not seen in the in vivo data, despite matched spatial resolution. This difference could relate to tissue changes due to death and pathology, but as yet we do not have sufficient data to determine the impact of pathology and death-related processes on these observations. Further data collected for the post-mortem dHCP study will provide insight.

Quantitative linear discriminant analyses separated limbic tracts from both the association and projection tracts in the in vivo and post-mortem data using either DKI or NODDI model parameters (Fig. 8). However, the subsequent subdivision of the isocortical tracts into projection and association tracts is less well defined in both datasets.This may be due to the structural and developmental difference between the limbic and the projection and association tracts: limbic tracts connect allocortical structures, have relatively short growth periods, and are arranged into well-defined bundles; by comparison, projection and association tracts connect isocortical structures, have longer growth periods, and are arranged in sagittal strata (Vasung et al., 2010).

For the DKI model parameters, mean diffusivity values are lower globally in the post-mortem data, which was only partially accounted for by tissue temperature differences. Additionally, mean kurtosis values are higher globally in the post-mortem data. Together these differences suggest greater microstructural complexity and reduced diffusivity in the post-mortem data, which could potentially be due to cell swelling effects during the peri- and post-mortem period. Such changes are a direct consequence of the processes of cell death (D’Arcy, 2019) with the peak in intracellular volume expansion being delayed at reduced temperatures (Janaway et al., 2009). There are however many factors surrounding the physiological state of the infant prior to death, which will compound and accelerate these cell changes, such as infection and resultant cytokine release, degree of supplemental oxygen, and use of intravenous fluids (McAdams and Juul, 2012; Truttmann et al., 2020). The cell swelling interpretation is consistent with the NODDI model parameters, where intra- and extra-neurite fractions are inverted in the post-mortem and in vivo data (Fig. 7), as the extracellular space in the post-mortem data would be reduced as a result of cell swelling.

The MD local minimum and MK local maximum overlap the T2w hyperintesity, which occur approximately 1mm deep to the cortical plate, interpreted as the superficial subplate (Pogledic et al., 2020). This would be consistent with the known selective vulnerability of subplate neurons to hypoxia-ischemia (McQuillen et al., 2003), which could cause effects reflective of cellular swelling on MD and MK to be greatest in this region. Downsampling the post-mortem data to match the spatial resolution of the in vivo dHCP data does not remove the MD and MK peaks, further suggesting that this is influenced by post-mortem effects, as opposed to a simple spatial resolution effect.

Cell swelling has been reported to be associated with reduced diffusivity in the extracelluar space, which might be due to an increased tortuosity (Chen and Nicholson, 2000; Toorn et al., 1996) in the reduced extracellular space or increased membrane space that cause more water to bind electrostatically to the membranes and adjacent macromolecules (Jelescu et al., 2014). This suggests the higher orientation dispersion in the post-mortem data may be influenced by post-mortem cell swelling.

Digging deeper into these initial promising findings and tackling these interesting challenges is an ongoing endeavour for the FMN study. Due to the successful implementation of the care pathway for this infant post-mortem research study, which was well received by the clinical team, research imaging centre, and the parents participating, further data collection is ongoing. We hope collaborative research on this open dataset will accelerate advances in understanding of developing human brain as well as pathology and death during the term and preterm periods.

## Methods

### Participant recruitment and infant transport

#### Study Participants

Neonates who died whilst receiving inpatient care at the John Radcliffe Hospital (Oxford University Hospital NHS Trust) Newborn Care Unit were screened for study. Eligible neonates were born live between 23 and 44 weeks’ gestation, to mothers aged 16 years or more, and could be scanned within 48 hours of death. The clinical care team, independent of FMN, determined the appropriateness of approaching parents to discuss the trial on an individual basis. Infants were excluded from the study if they had non-MRI compatible metallic implants. Ethical approval was obtained from the South Central Oxford B NHS Research Ethics Committee (REC reference: 19/SC/0154), and research was conducted in accordance with standards set by Good Clinical Practice guidelines and the Declaration of Helsinki.

#### Recruitment

Families that were identified as eligible for inclusion and expressed an interest in the study by the clinical team were then approached by the lead FMN research nurse (F.A.) who gave a verbal explanation of the purpose and goal of the study and recruited the family into the study. Written information was also provided via a Parent Information Leaflet. Parents had the opportunity to ask questions and were given as much time as they needed within the 48-hour window to consider their participation in the study. Written informed consent was provided by parents via a Consent Form. Parents who did not speak English were only to be approached if an unrelated adult interpreter was available.

#### Transfer

After death, the infant’s body is transferred to a cold cot (Flexmort CuddleCot mattress temperature maintained around 9-13°C) on the Newborn Care Unit. Once parents had time to mourn, collect mementos, have visits from family members, and hold a blessing ceremony if desired, the lead FMN research nurse liaised with scan operators to coordinate scan timing and transport of the infant’s body to the MRI centre onsite. Scanning occurs overnight, beginning after normal working hours to facilitate a discrete transfer. Two research nurses transported the infant in a covered pram containing a cooling mat, maintaining the infant body temperature at approximately 8 °C. Parents were invited to accompany their infant during transfer, or could say goodbye on the ward. On arrival at the MRI centre, the infant was screened for non-MRI compatible objects, removed from their swaddling and had temperature monitoring probes placed on their forehead and underarm (see Fig 2). The infant was then swaddled for imaging: first in gauze, then wrapped in an MR-compatible cooling blanket, and then several outer layers of cotton cloth to prevent condensation from the cooling setup. The infant was then placed in a custom-designed acrylic cradle. Under the supervision of an experienced scan operator, the infant’s body was transferred to the scanner bed within the cradle. Throughout the overnight scanning, the research nurse and scan operator remained at the imaging centre, regularly monitoring the scanner and setup, including the infant’s temperature. After completion of scanning the following morning, the infant was transferred to the mortuary by the research nurse following the Newborn Care Unit guidelines.

### Data acquisition workflow

#### Hardware setup

##### RF coil and cradle

Infants were scanned on a Siemens Magnetom 7T MRI scanner with a custom built transmit/receive neonate head coil (RAPID Biomedical) for this study, which consists of two transmit channels and 16 receive channels. The coil has an inner dimensions of 117mm (minimum diameter) × 135mm (maximum diameter) × 114mm (length), which can fit most infants up to term gestation. In the event of scanning a larger infant, a Nova head coil with 32 receive channels and a single transmit channel can be used. In addition, a 7T-compatible acrylic cradle was custom-designed to optimally position the infant’s body within the RF coil and ensure minimal handling of the infant’s body. The cradle size is 500mm (length) by 180mm (width), with a narrowed head support section (50mm) that fits within the neonatal head coil. The height of the cradle is adjustable such that the infant’s head can be placed at the center of the coil regardless of the head size.

##### Cooling System

Maintaining a consistent and low temperature during scanning is crucial to avoid tissue damage and to maintain stable signal and diffusion properties. As described in Supplementary Information S1.1, we conducted extensive testing on appropriate phantom objects, identifying that RF deposition was particularly problematic, and required active cooling. This led to the development of an active cooling system for this study. The custom cooling system consists of a water cooling pad to swaddle the infant (Plastipad Infant, Cincinnati Sub-Zero) and a recirculating cooler (F250, JULABO GmbH), which can control the water temperature between −10 and 40 °C with a stability of ±0.5 °C. To avoid artefacts due to the circulating water, water is doped to achieve a 5mM solution of manganese chloride, which has a very short T2 (3-5ms) (Thangavel and Saritaş, 2017). The cooler is not MR compatible, requiring it to be housed in the scan operating room beyond the 5-Gauss line. The temperature of the infant’s body was monitored throughout scanning using two fibreoptic temperature probes (Neoptix), sited on the forehead and underarm. Temperature measurements were recorded on a Windows PC with 0.1Hz sampling rate. Water hoses for the cooler and fibre optic cables for temperature monitoring were run through the waveguide panel connecting the scanner room with the operator suite. Supplementary Fig. 1 and Supplementary Information S1.1 provide details about temperature measurement during a series of pilot experiments and FMN scans of the first two infants.

#### Image acquisition

##### Structural protocols

Structural scans were based on the adult human connectome project (Van Essen et al., 2012) rather than dHCP since the latter protocols were specifically designed to mitigate motion artefacts in vivo. A 3D MPRAGE (magnetisation prepared rapid gradient echo) sequence was used to acquire T1w data using the following parameters: TR=2440ms, TE =3.35ms, inversion time 1500ms, flip angle 9°, field of view 150mm × 150mm × 102mm, matrix size 384×384×256, 0.4mm isotropic resolution, bandwidth 330Hz/pixel, GRAPPA factor 2, echo spacing 6.7ms, 4 repetitions, total scan time of 34 minutes. A 3D SPACE (sampling perfection with application-optimized contrast using different angle evolutions) sequence (Mugler, 2014) was used to acquire T2w data using the parameters: TR=2300ms, TE=586 ms, field of view 150mm × 150mm × 96mm, matrix size 384×384×240, voxel size 0.4mm isotropic, bandwidth 250Hz/pixel, GRAPPA factor 2, echo spacing 6.72ms, 6 repetitions, total scan time 58 minutes.

##### dMRI protocols

The diffusion acquisition protocol used a 2D simultaneous multi-slice spin-echo sequence with readout-segmented echo-planar imaging and monopolar diffusion preparation (Frost et al., 2015; Porter and Heidemann, 2009). The FOV was set to 150 × 150 × 104 mm^3^, in order to accommodate the head of a 44-week gestation infant measuring on the 95th percentile (Bastiani et al., 2019). Matrix size 188 × 188 × 130, 0.8mm isotropic resolution, GRAPPA (Griswold et al., 2002) acceleration=2, multi-band acceleration=2, TR/TE=10.8s/113ms, 7 readout segments and 0.4ms echo spacing. 280 diffusion weighed volumes were acquired for three b values 3000, 6000 and 9000 s/mm^2^ with unique diffusion encoding directions (64, 88 and 128 directions, respectively). 21 b=0 volumes were also acquired, with 17 using the same phase encoding direction as the high-b-value data, and four with reversed phase encoding direction, to allow correction of susceptibility-induced image distortion (Andersson et al., 2003; Smith et al., 2004). The total acquisition time for dMRI data is just under 7 hours. In the Supplementary Information S2, we provide an overview of this process with justifications for key decisions made. A comparison between post-mortem and in vivo dMRI protocols is shown in Table 1.

**Table 1:** dMRI acquisition protocol comparison. dwSE = diffusion-weighted spin echo; TR = repetition time; TE = echo time; EPI = echo planar imaging; MB = multiband; GRAPPA = generalized autocalibrating partially parallel acquisitions

|  | Post-mortem | In vivo |
| --- | --- | --- |
| Diffusion preparation | dwSE | dwSE |
| Readout trajectory | 2D segmented (x7) EPI | 2D single-shot EPI |
| Acceleration | MB2 + GRAPPA2 | MB4 |
| Resolution (mm) | 0.8x0.8x0.8 | 1.5x1.5x1.5 |
| TR (ms) | 10800 | 3800 |
| TE (ms) | 113 | 90 |
| Number of diffusion volumes | 301 | 300 |
| b-values (s/mm<sup>2</sup>) | 0, 3000, 6000, 9000 | 0, 400, 1000, 2600 |
| Number of volumes/directions | 21, 64, 88, 128 | 20, 64, 88, 128 |
| Acquisition time | 6h 50min approx | 19min approx. |

dMRI images were reconstructed off-line in MATLAB (Mathworks, Natick, MA) from raw k space data. The split slice-GRAPPA algorithm (Cauley et al., 2014) was used to separate aliased slices, followed by GRAPPA reconstruction of segmented EPI data using similar pipeline as in previous work (Frost et al., 2015). This large volume of raw data (~300G) was incompatible with the reconstruction computer’s native storage, requiring data to be streamed from the scanner to an offline computer using the *iceNIH_RawSend* tool developed by Jacco A. de Zwart.

##### Comparison with in vivo ultrasound

Cranial ultrasound as shown in Fig. 3 were obtained from the infant three days prior to the infant’s death and approximately four days prior to obtaining the accompanying MRI images. The images were obtained on a Siemens Acuson S2000 portable scanner using a 10v4 probe on the paediatric ‘neo head’ setting. This 2D probe was set up using a frequency of 10Hz, the depth set to 7cm and the gain at 63%. All images were obtained via the anterior fontanelle. In Fig. 3, the top image is a coronal view at the level of the third ventricle and thalami, the middle image is a left parasaggittal view through the left lateral ventricle showing anterior and posterior horns, and the bottom image is another coronal view, this time at the level of the posterior horns of the lateral ventricles (with choroids).

### Image processing

The cortical surface was extracted from the structural T1w and T2w images based on the dHCP neonatal structural processing pipeline (Makropoulos et al., 2018) with modifications to address the strong bias field at 7T. Specifically, the T2w/T1w ratio image was used as input instead of T2w image to eliminate receive field bias, and the FSL tool *fsl_anat* was subsequently used with the *strongbias* option to correct the transmit field bias.

Diffusion MRI data were pre-processed using the *top-up* and *eddy* tools (Andersson and Sotiropoulos, 2016) to correct for susceptibility and eddy current induced distortions. We require a transformation from diffusion space to the age-matched template (Serag et al., 2012) to perform automated tractography. *FLIRT* (Jenkinson and Smith, 2001) was used to register the average b=9000s/mm^2^ image to the T2w structural with 6 degrees of freedom. A non-linear registration from T2w image to the age-matched template was also performed with ANTs SyN (Avants et al., 2008). The linear transformation matrix was then combined with the non-linear field to produce the transformation needed in the tractography pipeline.

To investigate the effects of spatial resolution, low-resolution post-mortem T2w and dMRI data were generated by undersampling the original high-resolution data to 0.8mm and 1.5mm isotropic resolution, respectively, using *FLIRT* with trilinear interpolation. T2w structural and dMRI images of five age matched dHCP infants were used for comparison. Detailed comparison between post-mortem subjects and in vivo subjects is included in Supplementary Table S2.

### dMRI model fitting

After pre-processing, we analysed the dMRI data using a combination of approaches to extract microstructurally-relevant properties.

#### Diffusion kurtosis model

The diffusion kurtosis imaging (DKI) model (Jensen et al., 2005) is a higher-order extension of the diffusion tensor model. DKI fits data at multiple b-values to a second-order expansion to capture water diffusion that deviates from a Gaussian displacement distribution. DKI model parameters include the conventional diffusion tensor metrics (Basser et al., 1994) that can be estimated from data at a single b-value. Here we derived fractional anisotropy and mean diffusivity maps, as well as a map of principal diffusion direction from the eigenvector of the tensor. In addition, we estimated maps of mean kurtosis, which reflects the degrees of tissue heterogeneity. The DKI model was fit to the data using FSL’s *dtifit* command.

#### NODDI model

Data were also fit with the neurite orientation dispersion and density imaging (NODDI) model (Zhang et al., 2012). Whereas DKI aims to capture features of the signal, NODDI is a biophysical model that aims to estimate properties of different microscopic tissue compartments. NODDI attributes the dMRI signal to tissue compartments characterised by diffusion that is restricted (interpreted as intra-neurite spaces), hindered (extra-neurite) and unrestricted isotropic (CSF). The model also estimates an orientation dispersion index to account for fanning or crossing of neurite populations. In order to model stationary water trapped in small compartments or bound to membranes and other subcellular structures following death, an isotropic restricted compartment was also added, which is common for post-mortem tissue (Alexander et al., 2010). NODDI was fit using the GPU-based *cudimot* tool, which reduces the otherwise long computational times for NODDI fitting (Hernández et al., 2013).

#### Tractography

Diffusion data were fitted using a ball-and-stick model (Behrens et al., 2007; Jbabdi et al., 2012), which provides fibre orientation estimation for multiple fibre populations from each voxel. This model is capable of delineating unnecessary fibre populations by automatic relevance determination priors (Behrens et al., 2007). Tractography was then performed with the “baby autoPtx” method, a neonate-specific automatic tractography pipeline developed for dHCP dMRI processing (Bastiani et al., 2019), to dissect 16 white matter tracts including projection, association, callosal, cerebellar and limbic fibres. These segmentated tract masks were used for tract specific microstructure analysis.

### Tissue-specific dMRI analyses

#### Cortical plate radiality

Analysis of the developing cortical gray matter focused on the orientation of the primary diffusion direction in the middle of the extracted gray matter masks. Cortical radiality was calculated as the dot product of the principal eigenvector derived from diffusion tensor fitting and the surface normal vector (McNab et al., 2013). The calculation was performed using the *Connectome Workbench* tool (Marcus et al., 2011) and the surface reconstruction from the dHCP structural analysis pipeline. The principal eigenvector map derived in the diffusion space was projected to the midthickness surface of the cortical plate followed by radiality calculation with the normal vector on the same surface. The visualization of the surface rendering was performed with the Workbench*’s wb_view* tool.

#### Subplate microstructure

Regions-of-interest (ROIs) for the gray and white matter in the right hemisphere were generated using the dHCP structural pipeline segmentation outputs. The white matter ROI was fed into FSL’s *distancemap* command to generate a voxelwise map of the distance from the nearest gray-white matter boundary. This was used to characterise diffusion parameters as a function of distance value. In addition, diffusion parameters were averaged over the cortical plate and taken to be the value at zero distance. As distance values are discretised, the median parameter value was taken for a given distance value, and a smoothing spline curve fit to the data using MATLAB fit function (model type = smoothing spline, smoothing parameter = 0.99). The group median for the in vivo dHCP data was calculated as the median parameter value across all subjects for a given distance.

#### White matter microstructure

Of the 16 white matter tracts generated using the *baby autoPtx* protocols, analysis was limited to tracts fully localised in the right hemisphere to reduce the influence of pathological tissue. We classified these tracts into three categories (projection, limbic, and association) using the system that was adopted in the dHCP dMRI analysis pipeline (Bastiani et al., 2019). This resulted in 11 tracts, including three association (inferior longitudinal fasciculus, superior longitudinal fasciculus, uncinate fasciculus), three limbic (fornix, cingulate cingulum bundle, parahippocampal cingulum bundle) and five projection pathways (acoustic radiations, corticospinal tract, anterior thalamic radiations, posterior thalamic radiations, superior thalamic radiations).

To generate each white matter tract ROI, the tract’s voxelwise normalised tract density was thresholded at 0.001 and binarized. Cortical voxels were excluded using the gray matter mask generated during preprocessing. For each resultant tract ROI, the median dMRI parameter value was extracted for each DKI and NODDI parameter.

The Fisher’s linear discriminant ratio (LDR) was used to evaluate the separability between tract categories, which is calculated as the ratio between the inter-class and intra-class variances. Data were projected along the direction of the optimal linear discriminant vector that maximize the class-separability, and LDR was calculated from the projected data. Permutation tests were used to determine whether the clustering results performed signifantly better than chance: p-values were calculated by generating a LDR null distribution from 100,000 random permutations of tract categories and computing the fraction of permutation trials that give at least as high as the LDR obtained from the unpermuted tract categorization.

## Data availability

Source data in the paper are provided. Due to ethical restrictions, it is appropriate to monitor access and usage of the data as it includes highly sensitive information. Data sharing requests should be directed to. The dHCP data (Second Data Release) that used in this work are available online (http://www.developingconnectome.org/second-data-release).

## Code availability

FSL tools for dMRI processing and analysis are available via FSL (https://fsl.fmrib.ox.ac.uk/fsl/fslwiki). FSL Eddy tool for dMRI data preprocessing are available via https://fsl.fmrib.ox.ac.uk/fsl/fslwiki/eddy. DKI fitting was performed with FSL’s dtifit tool (https://fsl.fmrib.ox.ac.uk/fsl/fslwiki/FDT). Bias field correction was performed with FSL’s *fsl_anat* tool (https://fsl.fmrib.ox.ac.uk/fsl/fslwiki/fsl_anat). Linear image registration was performed with FSL *FLIRT* (https://fsl.fmrib.ox.ac.uk/fsl/fslwiki/FLIRT). Cudimot was used for NODDI fitting (https://users.fmrib.ox.ac.uk/~moisesf/cudimot/index.html). Tractography was performed with Baby Autoptx tool developed for the dHCP (https://doi.org/10.1016/j.neuroimage.2018.05.064). Processing pipeline for T2w structural data is available online (https://github.com/BioMedIA/dhcp-structural-pipeline). Connectome workbench tool can be found from HCP website (https://www.humanconnectome.org/software/connectome-workbench)

## Acknowledgements

The research leading to these results has received funding from the European Research Council under the European Union Seventh Framework Programme (FP/2007-2013)/ERC Grant Agreement no. 319456. We are grateful to the families who generously supported this trial. The Wellcome Centre for Integrative Neuroimaging is supported by core funding from the Wellcome Trust (203139/Z/16/Z). WW is supported by the Royal Academy of Engineering (RF\201819\18\92). KM is supported by the Wellcome Trust (WT202788/Z/16/A). LB, FA, REF, VM, FM, and RS are supported by the Wellcome Trust (207457/Z/17/Z). J.S. is supported by a IDEXLYON “IMPULSION 2020 grant (IDEX/IMP/2020/14) and the Labex CORTEX ANR-11-LABX-0042 of Université de Lyon. AFDH is funded by the Engineering and Physical Sciences Research Council (EPSRC, EP/L016052/1) and Medical Research Council (MRC, grant MR/L009013/1).

## Competing interests

The authors declare no competing interests.

## Notes

### Competing Interest Statement

The authors have declared no competing interest.

