## Supplementary for "The Forget-Me-Not dHCP study: 7 Tesla high resolution diffusion imaging in the unfixed post-mortem neonatal brain"

**Wu et al.**

### Supplementary Methods

#### S1: Active cooling system

##### S1.1 Temperature experiments

Substantial temperature rise during a scan has detrimental consequences to tissue, such as active damage or acceleration of degeneration (e.g., through autolysis), and to data, due to temporally varying water diffusivity and tissue relaxation properties. We hypothesised that the temperature changes would be more significant in the FMN study than previous post-mortem studies due to the use of high-power RF pulse for MB excitation, ultra-high field and long scan time.

RF energy transmitted by the excitation pulses is absorbed by the body and turned into heat within the tissue. For in vivo scanning, much of the RF-induced heating is dissipated by thermoregulatory mechanisms (blood vessel dilation that leads to increased blood flow), which is critical for preventing substantial temperature elevation. In post-mortem MRI scans, the body lacks thermoregulatory mechanisms, and as a result RF induced heat is more likely to accumulate in the body and lead to substantial temperature increase during the scan. Critically, substantial temperature increases might be expected to be incurred even for protocols that would be well within normal safety limits in vivo (i.e., < 1°C according to International Electrotechnical Commission 60601-2-23:2011).

In addition, temperature difference between the body and ambient air can also affect body temperature through heat exchange between skin and air. Because post-mortem infant temperature should be maintained at temperatures matching mortuary conditions (using a Flexmort CuddleCot mattress temperature maintained around 9-13°C while on the neonatal ward) in order to provide bereaved parents the option of spending time with their baby before transport to the mortuary, body temperatures of FMN infants were expected to be significantly lower than the ambient temperature (21°C) in the scanner room, which would cause temperature to rise independent of scanning. Moreover, the use of dMRI protocols with high b-value over long scan times are expected to increase ambient temperatures due to heavy gradient duty cycle.

To test our hypothesis, we scanned an unfixed porcine brain using a candidate diffusion protocol with the highest b value of 4000 s/mm$^2$ and measured tissue temperature continuously during the scan. The brain was extracted on the day of sacrifice and packed in a cylinder filled with fluorinert solution. The temperature was monitored using two fibreoptic temperature probes (Neoptix). One temperature probe was inserted into the brain, the other was placed at the end of scanner table to measure the ambient temperature. During a 6-hour diffusion scan, the tissue temperature measured inside the brain increased 5 °C, while the ambient temperature increased 2 °C (Fig. S1a). This would be consistent with both ambient heating due to gradient hardware heating (measured by the external probe) and additional heating due to RF energy deposition (reflected in the additional temperature rise in the internal probe). A conclusion from this experiment was that an active system was required to keep infants cool to avoid tissue damage, but also to stabilise temperatures to avoid unwanted confounds due to varying MR signal properties.

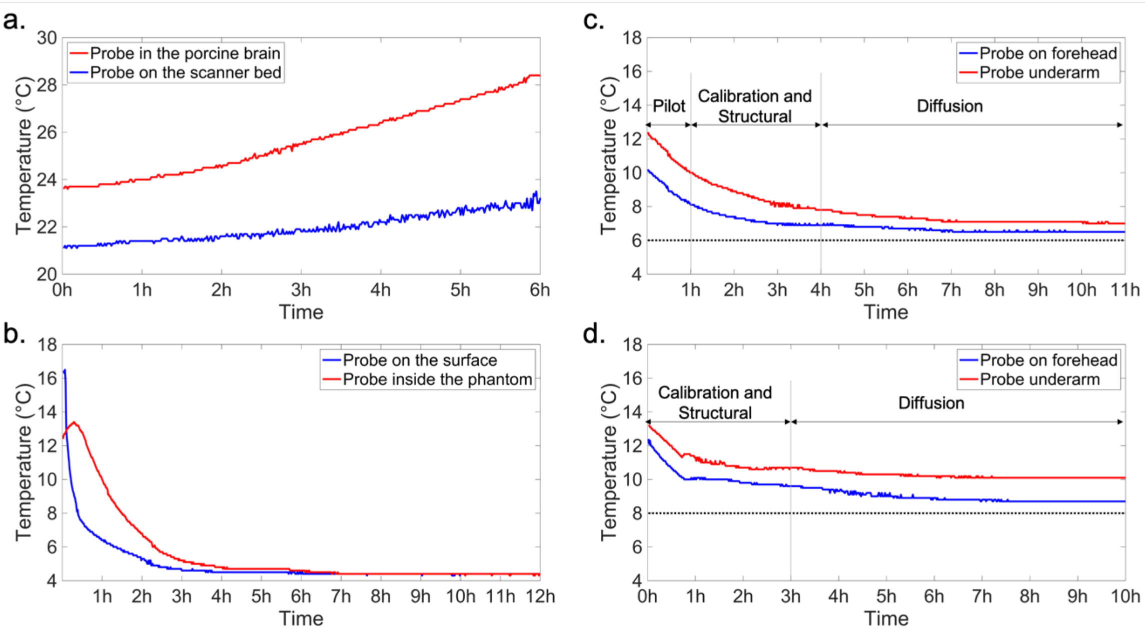

***Supplementary Figure 1. Temperature measured during different experiments and active cooling was applied during data acquisitions for b) – d).*** *a) Temperature of brain tissue (red) and ambient (blue) during a six-hour diffusion scan with a porcine brain. b) Temperature measured from the inside (red) and surface (blue) of a tissue-mimicking phantom. c) temperature measured during the first FMN scan. d) temperature measured during the second FMN scan. Two probes were placed on the forehead (blue) and underarm (red) of the infants during the FMN scans. The target temperatures set for the cooler in two FMN scans are shown as dotted horizontal lines.*

#### S1.2 Cooling system

We considered both gas- and water-based cooling approaches.

Gas cooling (circulating cold air) was initially expected to be preferable because it does not produce any signal in the image that might be confounded with the tissue signal. We built a prototype gas-cooling system fed from compressed air cylinders. This system was tested on a cylindrical phantom made from ground beef, but we found that a very high flow rate was needed in order to effectively cool the phantom. Maintaining such high flow rate for many hours was not considered to be practical or cost effective.

Our final design was thus based on water cooling, as described in detail in the main manuscript. The system is driven by a recirculating cooler (F250, JULABO GmbH) with a set target temperature that drives cold water through PVC tubing, which was divided into three sections. The first section is permanently connected to the cooler and the third section is permanently connected to the cooling pad that is wrapped around the infant. The middle section connects to the other two using non-spill valve hose couplings to prevent leakage of fluid while disconnecting. The use of two valves means that the sections connected directly to the cooler and blanket can be no longer than necessary, while the middle section provides most of the length needed to span between the waveguide and scanner isocentre (Fig. 2). With this setup, the first section of tubing remains permanently in place and setting up the cooling circuit simply requires connecting the two sets of valves, which takes less than one minute. The cooler works in a temperature range of -10 to +40 °C and can achieve a stability of ±0.5 °C. The total cost of the cooling system is about $3700.

To evaluate the efficacy of the final cooling system, we scanned a tissue mimicking phantom made from 500g of ground pork wrapped in the cooling blanket. We measured the temperature continuously during the scan, which included a five-hour structural scan followed by a seven-hour diffusion scan. One temperature probe was attached to the surface of the phantom and the other was inserted into the phantom. This experiment aimed to provide a conservative test of the capabilities of the cooling system. The phantom had been stored in a refrigerator prior to the experiment with a temperature of 12.4°C, which was thus below the room temperature, but well above the cooler's target temperature of 4°C. The temperature inside of the phantom increased by 1°C in the first half an hour (consistent with RF heating) and then decreased from 13.4°C to 5°C over 3 hours (Fig. S1b). The temperature on the phantom surface close to the cooling blankets decreased immediately, asymptotically approaching 5°C in 2 hours. The results demonstrate the cooling system can both counteract sources of heating within a short time and maintain a stable working temperature over a sustained scanning protocol.

The cooling system was used in the two FMN scans. The infants were wrapped in three layers (Fig. 2). The innermost layer was gauze, which was used to protect the skin from contact with the cooling blanket. The middle layer was the cooling blanket. The outer layer was cotton cloth used to block airflow, which might cause condensation on the surface of the cooling blanket. Two fibre optic temperature probes were placed on the forehead and underarm, respectively. In the first FMN scan, the target temperature was set to 6°C. As shown in Fig. S1c, the temperature measured at the underarm decreases from 11.9°C to 8°C over three hours, and slowly decreases by no more than 1°C over the following 7-hour diffusion scan, with a final temperature of 7°C. The temperature at the forehead decreases from 9.8°C to 6.9°C over 3 hours and decreases by no more than 0.5 °C over the following 7 hours. In the second FMN scan, the target temperature was set to 8°C. As shown in Fig. S1d, the temperature measured from the underarm decreases from 13.2°C to 10.7°C over 3 hours, and then decreases by no more than 1°C over the following 7-hour diffusion scan with a final temperature of 10.1°C. The temperature at the forehead decreases from 12.4°C to 9.6°C over 3 hours and decreases by no more than 1 °C over the following 7 hours with a final temperature of 8.7°C. Note the temperature set on the cooler is the target temperature of fluid in the cooler tank, which will be warmed up when reaching the cooling pad due to heat exchange with the tubing. Therefore, the final temperature measured from the probes was slightly higher than the target temperature set on the cooler.

### S2: Optimisation of diffusion acquisition

##### S2.1 Diffusion preparation scheme

Post-mortem diffusion imaging in whole human brains has been dominated by two diffusion preparation schemes: diffusion weighted spin-echo (DW-SE) (Miller et al., 2011, 2012) and diffusion weighted steady-state free precession (DW-SSFP) (McNab et al., 2009; Miller et al., 2012). DW-SE is the workhorse for most diffusion MRI studies, including the in vivo dHCP study. However, DW-SE suffers from a trade-off between diffusion contrast (requiring long TE to achieve high b values) and SNR (requiring short TE to reduce T2 decay). This is a problem for post-mortem imaging of fixed, adult brains when tissue T2 and ADC are both low (Miller et al., 2012; Roebroeck et al., 2019). However, in the infant brain, tissue T2 is characteristically longer than adult brain due to higher water content (Leppert et al., 2009), which might allow a longer TE without substantially compromising SNR. DW-SSFP has been demonstrated to provide benefits for post-mortem diffusion imaging due to its ability to overcome the trade-off between contrast and SNR, and thus achieve high SNR efficiency in tissue with short T2 (McNab et al., 2009; Miller et al., 2012). One downside of DW-SSFP is that signal formation mechanisms are different from DW-SE, meaning that analysis methods developed for DW-SE need to be adapted for DW-SSFP data (McNab and Miller, 2010).

First, we describe our approach to predicting T2 in unfixed, post-mortem infant brain at 7T. The predicted benefits of different diffusion preparation scheme were calculated from estimated tissue T2 and ADC for unfixed post-mortem infant brain. Previous work by (Thayyil et al., 2012) reported T2 values of 280ms (WM) and 210ms (GM) from unfixed post-mortem infant brain at 1.5 T. Using a linear fit of field-dependent T2 changes based on literature data (Deistung et al., 2008; Uludağ et al., 2009), we predicted T2 values of 178ms (WM) and 121ms (GM) for unfixed infant brains at 7T. Since this prediction is based on a study with very similar body temperature (8°C), the temperature related T2 difference was assumed to be negligible.

In addition, we needed an approximate ADC for unfixed, post-mortem infant brain at low temperature. Two previous studies have reported tissue ADC values from unfixed post-mortem infants, finding ADCs of 0.2-0.3 $mm^2/ms$ at 4-8°C (Papadopoulou et al., 2016; McDowell et al., 2018). The low ADC values reported in these two studies are driven in part by the temperature dependency of water diffusivity. As described above, FMN uses active cooling to keep the infant's body temperature around 5-10°C, resulting in low ADC that necessitate high b-value to achieve good diffusion contrast. In our optimisation, we assumed ADC values to be similar to previous findings.

Based on these estimates of T2 and ADC, we are able to predict signal levels for DW-SE and compare them to known protocols. The HCP (human connectome project) 7T diffusion protocol for $b=2000s/mm^2$ requires TE=71ms, which is applied to adult brain with ADC≈1 mm$^2$/ms (i.e. a factor of ~5 higher than predicted for FMN infants). In order to achieve equivalent contrast in FMN, we would need to aim for b=10,000 s/mm$^2$, which on our system requires TE=120ms. Assuming T2=178ms for FMN and T2=45ms (Uğurbil et al., 2013) for HCP, the raw signal level of WM for FMN is predicted to be 2.5X higher than HCP. Given the HCP 7T protocol uses 1.05mm isotropic resolution with single-shot acquisition and FMN uses 0.8mm isotropic resolution with segmented acquisition (7 segments), the SNR of FMN data will be about ~3X higher than HCP.

This theoretical prediction suggests DW-SE is a viable option for the FMN study. Given that DW-SE also provides consistency with the in vivo dHCP study and compatibility with the majority of current analysis pipelines, we elected to use DW-SE preparation for the FMN diffusion protocol.

**S2.2 Q-space sampling**

The first consideration regarding q-space sampling was how to distribute data in q-space for our target scan time. We aligned our sampling scheme to the in vivo dHCP, with 64, 88 and 128 directions for low, medium and high b values, respectively (Tournier et al., 2020). This sampling aims to achieve a balance of dense angular coverage in a given shell to resolve fibre orientations for robust tractography, while also covering three b shells to allow more advanced modelling of diffusion MRI signal (Jensen et al., 2005; Zhang et al., 2012). Matching the in vivo dHCP q-space sampling scheme helps to harmonise FMN to this larger study.

A less straightforward consideration is the specific b-values for the three shells. The in vivo dHCP study acquires data at b=400,1000,2600 s/mm$^2$ (Bastiani et al., 2019). The FMN study requires higher b values due to the reduction of diffusivities as a result of lower tissue temperature and death, which previous studies predict to be ~5-fold lower compared to in vivo (0.2-0.3) (Papadopoulou et al., 2016; McDowell et al., 2018). Given the sparse literature on these diffusivities, we decided to prepare three sets of diffusion protocols before the first scan, with an upper b value of 4000, 6000 and 9000 mm$^2$/s. These protocols were chosen to enable us to rapidly pilot under conditions where SNR versus contrast is the limiting factor (favouring lower or higher b-values, respectively). The lowest b-values would provide reduced contrast relative to the in vivo dHCP protocol even if only accounting for temperature-induced reductions in diffusivity, while b=9000 s/mm$^2$ approaches the maximum b-value that is allowed in the current sequence (b_max=10000 s/mm$^2$). Other protocol parameters (e.g., TE, TR) were optimized for each protocol in accordance with the b value.

Due to the sensitive nature of recruitment in FMN, we did not feel it appropriate to dedicate any session to pilot scanning, requiring a more rapid and flexible approach to evaluating different b-value options. During the first FMN scan, we allocated one hour at the beginning of the scan session to acquire and reconstruct diffusion datasets with b=4000, 6000 and 9000 s/mm$^2$. From this rapid turnaround pilot scan, we found the ADC of brain tissue in the first FMN infant was ~0.2 mm$^2$/ms, consistent with previous findings (Papadopoulou et al., 2016;

McDowell et al., 2018). Diffusion MRI data acquired with b=9000 s/mm$^2$ provided considerably improved image contrast than those with lower b values, while still preserving image SNR. We therefore decided to use the protocols with an upper b value of b=9000 s/mm$^2$, with the other two shells at b=3000 and 6000 s/mm$^2$.

Long scan-time diffusion protocols place a high demand on the gradient system, with each individual scan inducing thermal heating that increases with b-value. If this heating exceeds the capacity of the scanner's cooling system, the scanner will shut down to avoid damage. To minimise heating for our given set of directions and b-values, the three b shells were segmented into subsets and interleaved (with four, five and eight subsets for b=3000, 6000 and 9000 s/mm$^2$, respectively). In the scan, the 9 sub-groups of b=3000 s/mm$^2$ and b=6000 s/mm$^2$ protocols were interleaved with the 8 sub-groups of b=9000 s/mm$^2$ protocols, in order to avoid running gradient coils at high duty-cycle continuously for a long period of time. As a worst scenario check, we also tested the b=9000s/mm$^2$ protocol in a continuous run of 3 hours on a phantom, which completed without gradient overheating.

In addition to the 280 diffusion-weighted volumes, 21 b=0 volumes were also acquired including 17 with the same phase-encode direction as the diffusion-weighted data, and 4 with reversed phase-encode direction, the latter for use in distortion correction (Andersson et al., 2003; Smith et al., 2004). The b=0 volumes were acquired at the beginning of each sub-set and thus have a relatively even distribution across the scan. This "timeseries" of b=0 volumes were used to monitor potential detrimental effects like signal drift.

##### S2.3 Acceleration and scan time

The primary considerations for acceleration are scan time, dominated by the simultaneous multi-slice (SMS) acceleration, and image quality, reflecting both slice-wise and in-plane acceleration. We consider the former in this section and the latter in the next section (S2.4).

In SMS, multiband (MB) RF pulses are used to excite multiple brain slices simultaneously. These slices then have shared diffusion encoding and readout, enabling the repetition time to be reduced. SMS with acceleration factor of MB=N provides a ~N-fold scan time reduction. Signals from individual slices are subsequently separated using unaliasing algorithms with coil sensitivity profile measured from a separate scan (Cauley et al., 2014).

A major issue with SMS excitation is the considerably higher power deposition by the radiofrequency (RF) pulses, leading to sample heating. This effect is usually measured by means of 'specific energy absorption rate' (SAR) that is related to the average energy dissipated in the body tissue per unit of mass and time. Modern MRI scanners monitor SAR in real time and do not allow sequences to run if SAR exceeds the limit. Our 7T scanner incorporates two SAR modes: a 'burst' mode, which allowed the sequence to run at the full SAR limit (100%) continuously for a maximum of ~6 minutes, and a 'conservative' mode, which allowed the sequence to run at ~80% of the full SAR limit continuously for long periods. Due to our longer scans, the FMN study needs to use the ‘conservative’ mode, leading to a stricter SAR constraint.

For our scanner’s specific implementation of diffusion-weighted spin echo with readout-segmented EPI, we compared different combinations of MB and in-plane phase encoding acceleration factor (MB=1,2,3,4 and R=1,2,3,4). We then explored the achievable parameter space of MB factors to determine the optimal choice by evaluating SAR compatibility and the number of diffusion volumes that can be achieved. Acceleration factors higher than 4 are not compatible with the post-mortem neonate coil and were not considered.

A first consideration is whether the scanner will impose SAR restrictions for a given protocol, which would disallow the protocol from being run (note that this is a separate consideration from the tissue heating effects discussed at length above). Fig. S2a shows scanner predicted SAR for different acceleration protocols with minimum allowable TRs. High MB factors are associated with higher SARs. Most MB3 and MB4 protocols with b=4000s/mm$^2$ and minimum allowable TRs exceed the conservative SAR limit. The b=9000s/mm$^2$ protocols result in slightly lower SAR than the b=4000s/mm$^2$ protocols, due to the longer TR required for higher b-value; while accelerations with MB<4 are generally not SAR limited for b=9000s/mm$^2$, the scanner will impose SAR restrictions at MB=4. We implemented reduced-SAR MultiPINS RF pulses (Eichner et al., 2014) (Fig. S2a). These pulses generally enable all combinations of acceleration factors to be run without encountering SAR restrictions. However, MultiPINS pulse incurs off-resonance sensitivity due to the longer pulse duration (Eichner et al., 2014). A simpler approach is to simply increase the TR beyond the minimum. Figure S2b shows minimum allowable TRs for all protocols, where an increase in TR is needed (dashed line) in some protocols with MB=3,4 to be SAR compatible.

For protocols that will run without SAR restriction, a second consideration is the total number of image volumes that is achievable within the 7-hour scan time limit. Fig. S2c demonstrates this dependence, which leads to the exclusion of all MB=1 protocols and MB2R1 protocol, which cannot provide at least 301 image volumes in 7 hours.

The result of these evaluations of multi-band factors relating to scan time considerations are that: (i) MB>1 is needed to achieve our desired q-space sampling and (ii) MB>2 would require some strategy to avoid SAR restrictions (altered RF pulses or non-optimal TR).

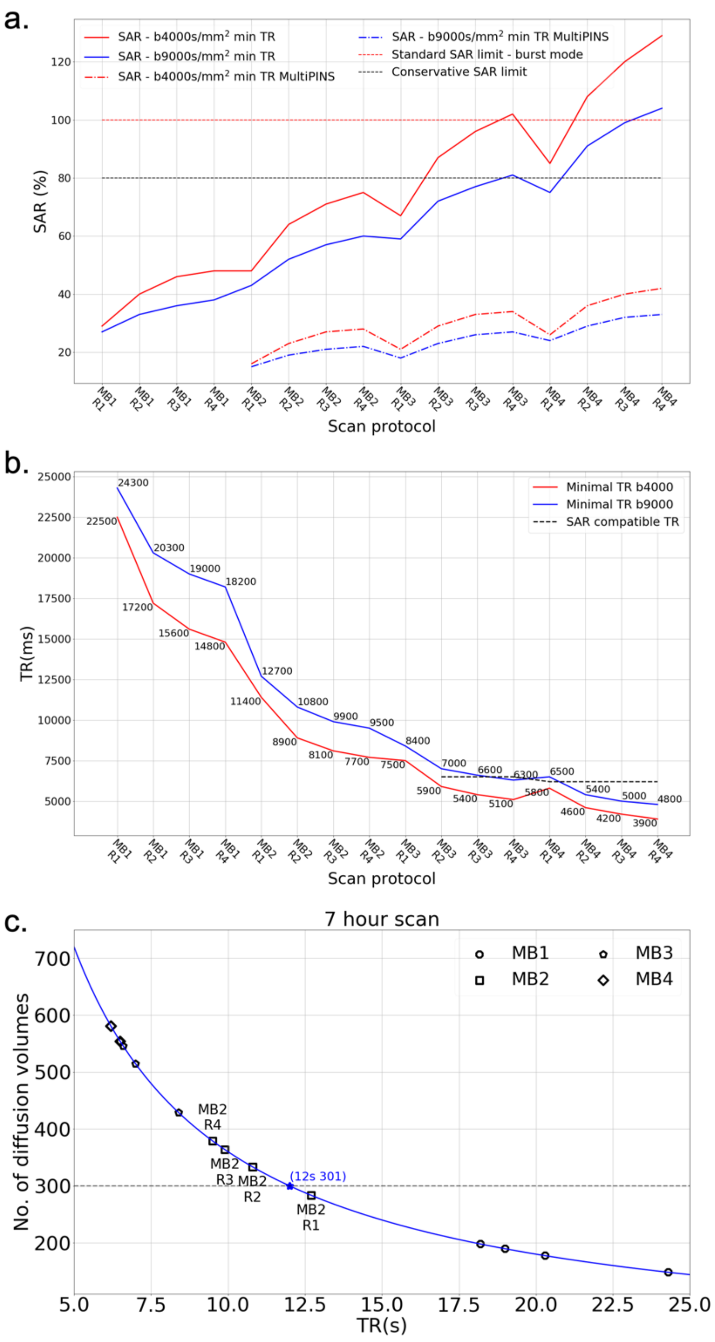

***Supplementary Figure 2. Simulation of accelerations for different protocols*** *a) Scanner predicted SAR for different acceleration protocols with minimum allowable TRs. In order to run a protocol consecutively for long hours, sequence SAR needs to be lower than the conservative SAR limit (black dot line). Burst mode allows a sequence to run at full SAR limit for a maximum of 6min (red dot line). Compared to conventional Multiband pulses (solid lines), MultiPINS pulses are less SAR sensitive (dot dash lines). Protocols with b=9000s/mm² (blue solid line) have a lower SAR than the b=4000s/mm² protocols (red solid line) due to the long TRs required for large b-values. b) Minimum allowable TRs for all protocols under investigation. To be SAR compatible, some protocols with high MB factors need to use a higher TR (dash line). c) The relation between the achievable number of diffusion volumes*

*with TRs. In order to acquire at least 301 image volumes in the 7-hour scan, the TR should not exceed 12s (dash line), which excludes five protocols including all MB=1 protocols and MB2R1 protocol.*

##### S2.4 Acceleration and image fidelity

In addition to considerations relating to scan time discussed in S2.3, acceleration decisions also impact on image artefacts. In-plane acceleration reduces blurring and distortion, while both slice-wise and in-plane acceleration lead to residual aliasing artefacts and noise amplification. As described in the following section S2.5, our primary tool for reducing blurring and distortion is readout-segmented EPI; however, this comes at a cost of scan time. Hence, we also considered acceleration along the phase encoding direction, which does not increase scan times but does incur a factor of $\sqrt{R}$ SNR loss for an R-fold acceleration. We explored the achievable parameter space to determine the impact of different acceleration strategies on blurring, distortion and unaliasing artefacts.

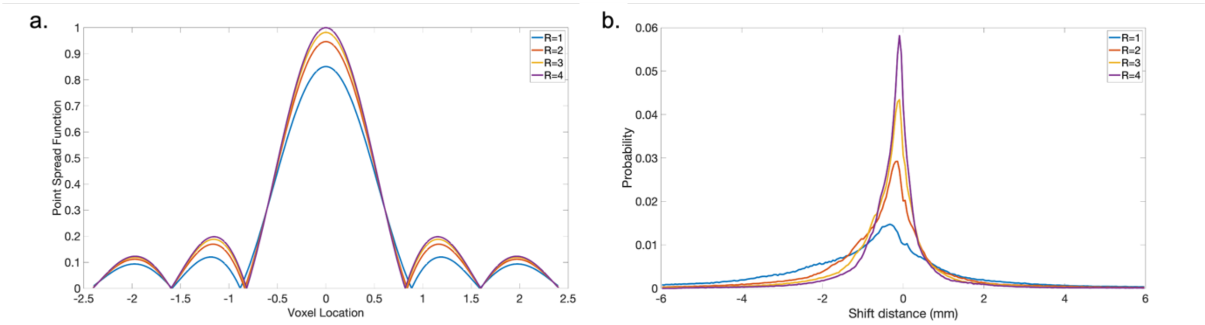

***Supplementary Figure 3. Simulation of blurring and distortion.*** *a) Blurring simulation: plot of voxel point spread functions along phase encoding direction for different phase encoding acceleration factors with an estimated T2* of 83ms. b) Distortion simulation: histogram of voxel shift distances simulated based on a representative B0 field map that was acquired at 3T from a healthy infant and linearly scaled to 7T.*

| **Phase encode acceleration** | **FWHM (voxel)** | **90$^{th}$ percentile shift distance (mm)** |
|---|---|---|
| R=1 | 1.29 | 2.16 |
| R=2 | 1.24 | 1.08 |
| R=3 | 1.23 | 0.72 |
| R=4 | 1.22 | 0.54 |

***Supplementary Table 1. Quantitative measurement of image blurring and distortion for different phase encoding acceleration factors****. Image blurring is evaluated with the full width at half maximum (FWHM). Distortion is assessed with the 90$^{th}$ percentile of the shift distance of all voxels in the brain.*

Fig. S3 shows a realistic simulation of blurring and distortion effects for different in-plane acceleration factors based on an estimate of WM T2* (83ms) for the FMN infant and a representative B0 field map that was acquired at 3T from a healthy infant and linearly scaled up to 7T. Blurring was measured using the full width at half maximum (FWHM) of the point spread function (Fig. S3a); distortion was assessed with the 90$^{th}$ percentile of the shift distance of all voxels in the brain (Fig. S3b). Based on these simulations, R=1 is expected to contain substantial distortion, with a large number of voxels shifting >2mm (3 or more voxels in our final target resolution). Using a small acceleration factor R=2 is expected to substantially reduce this artefact. The largest reduction is gained from R=1 to 2, while using higher acceleration factor only provides marginal improvement, particularly for blurring artefacts according to the FWHM measure (Table. S1).

We tested all protocols on a phantom and evaluated the quality of the reconstructed images. Fig. S4a shows results with different combinations of MB=2,3,4 and R=2,3,4. The normalised root mean square error (NRMSE) was calculated for each result using MB=1 image as a reference, which is shown on the bottom right corner of each image in Fig. S4a. As expected, image quality is better with a lower acceleration factor, and reconstruction breaks down when both SMS acceleration and phase encoding acceleration factors are high. The MB2R2 result has the lowest NRMSE, followed by MB3R2, MB2R3 and MB4R2 results. These results suggest that, while the product of the two acceleration factors is generally the overriding factor, in practice the reconstruction tolerates high MB factors better than high in-plane acceleration factors.

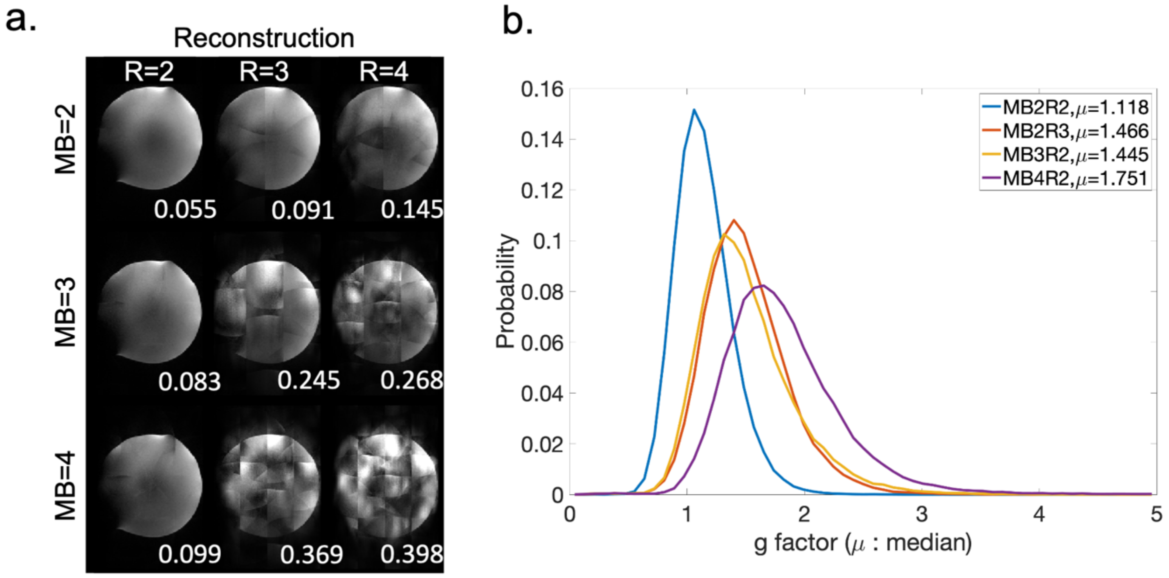

***Supplementary Figure 4. Evaluation of reconstruction performance for different acceleration protocols.*** *a) Reconstructed phantom images for different combinations of MB acceleration factors and phase encoding acceleration factors R.* Normalised root mean square error is shown on the bottom right corner of each image. *b) Histogram of g-factors over all voxels for MB2R2, MB3R2, MB4R2 and MB2R3. μ = distribution median.*

These four protocols are further compared for SNR penalty based on g-factor, which quantifies the reduction of SNR due to coil-geometry and sampling trajectory. Fig. S4b shows the histogram of g-factor over all voxels for these four protocols, where MB2R2 protocol outperforms the other three by a large margin and therefore incurs a much lower SNR penalty. The overall SNR efficiency (SNR per unit time) of MB2R2 protocol is ~10% higher than the SMS3R2 protocol, which is the second-best protocol.

The result of these evaluations of acceleration with respect to image quality are that: (i) R>1 will provide major improvements in blurring and distortion, and (ii) R>2 incurs major image artefacts. Hence, for in-plane accelerations R=2 is the most robust option. Taking this together with the considerations from S2.4 regarding MB factors, we made the decision to use MB=2 and R=2.

##### S2.5 Readout trajectory and spatial resolution

A major aim of the FMN study is to achieve higher resolution that cannot be easily achieved in vivo. The SNR required to achieve this aim is directly enabled by the use of ultra-high field and long scan times. However, the combination of ultra-high field and high resolution makes EPI-based acquisition more susceptible to image distortion and T2 blurring. Segmented EPI reduces image distortion and blurring at the cost of longer scan time to acquire multiple segments. There are two types of segmented EPI available: phase-encode segmentation (Butts et al., 1997; Atkinson et al., 2000) and readout segmentation (Porter and Heidemann, 2009). Since the primary distinction between these two approaches for dMRI relates to the use of navigator-based motion correction, they would both be suitable for post-mortem imaging. The FMN study opted to use readout-segmented EPI, which was available on our 7T platform as a "work-in-progress" package from the scanner vendor (Siemens Healthineers, WPI_591D). The sequence was modified to incorporate simultaneous multi-slice excitation and to remove motion navigator acquisition, which is not needed in post-mortem scanning. Removing the navigator acquisition reduced both SAR and scan time.

The decoupling of spatial resolution from the duration of the readout in readout-segmented EPI enables one to trade off scan time for reduced distortion and blurring; this flexibility also makes protocol optimisation more complicated than with single-shot EPI. Achieving high spatial resolution requires a large number of readout segments and slices. The scan time for each diffusion volume is dictated by the product of the number of slices and the number of readout segments. Scan times can then be reduced through the use of simultaneous multi-slice acceleration, which in effect reduces the time per slice, and in-plane acceleration, which enables the use of fewer segments for a given level of distortion.

Our protocol optimisation targeted the maximum resolution that could be achieved under constraints relating to (i) isotropic resolution, (ii) maximum scan time, (iii) achievable acceleration, and (iv) maximum tolerable distortion/blurring. In order to acquire 301 dMRI volumes (see section S2.2) within the allotted seven hours, the scan time per diffusion volume should not exceed 83s. As described in section 2.3 and 2.4, the optimal acceleration factors are R=2 (in-plane) and MB=2 (slice-wise). Under these constraints, the highest achievable resolution is 0.8mm, acquired using 7 readout segments.

| | Subject (session) | Sex | GA (weeks) | PMA (weeks) | PNA (days) | BW (grams) | Head motion (mm)** |
|---|---|---|---|---|---|---|---|
| Post-mortem | FMN infant | M | 23.0 | 29.6 | 46 | 545 | n/a |
| In vivo | CC00389XX19 (119100) | M | 28.7 | 29.9 | 8 | 825 | 2.89 |
| | CC00530XX11 (153600) | F | 26.14 | 29.29 | 22 | 960 | 3.30 |
| | CC00618XX16* (177201) | F | 27.43 | 29.86 | 17 | 760 | 7.47 |
| | CC00657XX14 (193700) | M | 28.14 | 29.86 | 12 | 1255 | 1.68 |
| | CC00672AN13 (197601) | M | 28.71 | 30 | 9 | 1310 | 1.77 |
| | CC00735XX18 (222201) | M | 25.57 | 29.29 | 26 | 990 | 1.28 |

***Supplementary Table 2: Subject comparison.*** *GA = gestational age (at birth); PMA = postmenstrual age (at scan); PNA = postnatal age (at scan); BW = birth weight. * = excluded from analysis because of data quality issues. For subject CC00618XX16 in the in vivo study, this is likely due to large head motions during the MRI scan. ** = average relative translational head motion estimated with FSL Eddy.*

Smith, S.M., Jenkinson, M., Woolrich, M.W., Beckmann, C.F., Behrens, T.E.J., Johansen-Berg, H., Bannister, P.R., De Luca, M., Drobnjak, I., Flitney, D.E., Niazy, R.K., Saunders, J., Vickers, J., Zhang, Y., De Stefano, N., Brady, J.M., Matthews, P.M.,

2004. Advances in functional and structural MR image analysis and implementation as FSL. Neuroimage 23 Suppl 1, S208-219. https://doi.org/10.1016/j.neuroimage.2004.07.051
Thayyil, S., De Vita, E., Sebire, N.J., Bainbridge, A., Thomas, D., Gunny, R., Chong, K., Lythgoe, M.F., Golay, X., Robertson, N.J., Cady, E.B., Taylor, A.M., 2012. Post-mortem cerebral magnetic resonance imaging T1 and T2 in fetuses, newborns and infants. European Journal of Radiology 81, e232–e238. https://doi.org/10.1016/j.ejrad.2011.01.105
Tournier, J.-D., Christiaens, D., Hutter, J., Price, A.N., Cordero-Grande, L., Hughes, E., Bastiani, M., Sotiropoulos, S.N., Smith, S.M., Rueckert, D., Counsell, S.J., Edwards, A.D., Hajnal, J.V., 2020. A data-driven approach to optimising the encoding for multi-shell diffusion MRI with application to neonatal imaging. NMR in Biomedicine 33, e4348. https://doi.org/10.1002/nbm.4348
Uğurbil, K., Xu, J., Auerbach, E.J., Moeller, S., Vu, A.T., Duarte-Carvajalino, J.M., Lenglet, C., Wu, X., Schmitter, S., Van de Moortele, P.F., Strupp, J., Sapiro, G., De Martino, F., Wang, D., Harel, N., Garwood, M., Chen, L., Feinberg, D.A., Smith, S.M., Miller, K.L., Sotiropoulos, S.N., Jbabdi, S., Andersson, J.L.R., Behrens, T.E.J., Glasser, M.F., Van Essen, D.C., Yacoub, E., 2013. Pushing spatial and temporal resolution for functional and diffusion MRI in the Human Connectome Project. NeuroImage 80, 80–104. https://doi.org/10.1016/j.neuroimage.2013.05.012
Uludağ, K., Müller-Bierl, B., Uğurbil, K., 2009. An integrative model for neuronal activity-induced signal changes for gradient and spin echo functional imaging. NeuroImage 48, 150–165. https://doi.org/10.1016/j.neuroimage.2009.05.051
Zhang, H., Schneider, T., Wheeler-Kingshott, C.A., Alexander, D.C., 2012. NODDI: Practical in vivo neurite orientation dispersion and density imaging of the human brain. NeuroImage 61, 1000–1016. https://doi.org/10.1016/j.neuroimage.2012.03.072
